# Over-optimism in unsupervised microbiome analysis: Insights from network learning and clustering

**DOI:** 10.1101/2022.06.24.497500

**Authors:** Theresa Ullmann, Stefanie Peschel, Philipp Finger, Christian L. Müller, Anne-Laure Boulesteix

## Abstract

In recent years, unsupervised analysis of microbiome data, such as microbial network analysis and clustering, has increased in popularity. Many new statistical and computational methods have been proposed for these tasks. This multiplicity of analysis strategies poses a challenge for researchers, who are often unsure which method(s) to use and might be tempted to try different methods on their dataset to look for the “best” ones. However, if only the best results are selectively reported, this may cause over-optimism: the “best” method is overly fitted to the specific dataset, and the results might be non-replicable on validation data. Such effects will ultimately hinder research progress. Yet so far, these topics have been given little attention in the context of unsupervised microbiome analysis. In our illustrative study, we aim to quantify over-optimism effects in this context. We model the approach of a hypothetical microbiome researcher who undertakes three unsupervised research tasks: clustering of bacterial genera, hub detection in microbial networks, and differential microbial network analysis. While these tasks are unsupervised, the researcher might still have certain expectations as to what constitutes interesting results. We translate these expectations into concrete evaluation criteria that the hypothetical researcher might want to optimize. We then randomly split an exemplary dataset from the American Gut Project into discovery and validation sets multiple times. For each research task, multiple method combinations (e.g., methods for data normalization, network generation, and/or clustering) are tried on the discovery data, and the combination that yields the best result according to the evaluation criterion is chosen. While the hypothetical researcher might only report this result, we also apply the “best” method combination to the validation dataset. The results are then compared between discovery and validation data. In all three research tasks, there are notable over-optimism effects; the results on the validation data set are worse compared to the discovery data, averaged over multiple random splits into discovery/validation data. Our study thus highlights the importance of validation and replication in microbiome analysis to obtain reliable results and demonstrates that the issue of over-optimism goes beyond the context of statistical testing and fishing for significance.

## 1 Introduction

The popularity of microbiome research has surged in recent decades. Many hypotheses about the human microbiome, as well as the microbiome of other species or in various environments, are postulated and tested each year. At the same time, new statistical and computational methods for analyzing microbiome data are continually introduced. Microbiome analysis has yielded exciting results, leading to high hopes for new treatment and prevention options in medicine [1, 2].

In such a fast-moving and promising research field, validation is of vital importance to ensure the reliability of new results. Yet such practices may sometimes be neglected in favor of chasing new hypotheses. There is a certain danger of *over-optimism* in the field: New and exciting results might turn out to be non-replicable, i.e., they cannot be confirmed in studies with independent data. While a discussion about validation and replication has emerged in microbiome research in recent years [3, 4], it is not as advanced as in other fields such as psychology, where the so-called “replication crisis” has received considerable attention [5]. There is a lack of studies which illustrate the validation process in microbiome analysis and quantify over-optimism and (non)replicability. In particular, scant attention has been given to these topics in relation to *unsupervised* microbiome data analysis, e.g., network analysis and clustering.

In the present paper, we take a step toward filling this gap. We illustrate how overoptimism can arise in unsupervised microbiome analysis using three unsupervised “research tasks” as examples: clustering bacterial genera, finding hubs in microbial networks, and differential network analysis. The underlying idea is to model the approach of a “hypothetical researcher” who has these research tasks in mind and is confronted with a variety of methods to choose from. Due to uncertainty about the appropriate method to apply in the present case, the researcher might be tempted to try different analysis strategies and pick the “optimal result” for each task. We quantify the over-optimistic bias that can arise out of choosing the “best” method in this way, by validating the optimized results on validation data (which we will define shortly). Our primary interest does not lie in any of the three specific research tasks, but rather in demonstrating the importance of validation and the necessity of avoiding questionable research practices. Through this illustrative study, we aim to raise awareness for these topics in microbiome analysis.

We now explain our usage of the terms “over-optimism”, “validation”, and “replication”. Broadly speaking, over-optimism may result from two sources of *multiplicity*: a) multiplicity of (tested) hypotheses or b) multiplicity of analysis strategies. It is well known that *multiple testing* (i.e., testing multiple hypotheses on a dataset) can lead to false-positive results due to the accumulation of the type I-error probability. Such problems may appear in microbiome research, e.g., when testing many associations of microbiome-related variables with health-related variables and only reporting the significant results [3]. However, even when considering only a single hypothesis, the *multiplicity of analysis strategies* [6] – which we focus on in this paper – may lead to varied results and the potential for selectively reporting only the best ones. Researchers must make several choices about their analysis strategy (a mechanism known as “researcher degrees of freedom”, [7]), including data preprocessing (e.g., normalization) and statistical analysis in a narrower sense. Often, multiple analysis strategies are possible and sensible, which leads to *method uncertainty* [8] because it is not necessarily clear which analysis choice is the best one. In microbiome analysis, for example, a large number of methods for estimating and analysing microbial association networks exists [9], from which the researcher must choose.

In such situations, there is a temptation for the researcher to try different methods and then pick the one that yields the best results. This approach might be considered sensible: Finding the “best” method for the data appears to be a natural goal. However, when the number of tried methods is high, there is a substantial danger of “overfitting” the analysis to the present dataset. The best-performing method might thus perform well on the data currently used, but perhaps not as well on a validation dataset due to sampling variability – in other words, the optimized result cannot be (fully) validated or replicated on the validation data. Here, we define “replication” as applying the same methods of a study to new data [4]; see [10] for a more extensive discussion of the concept of replication. “Validation”, as we use it, is a broader term: A result is reappraised on a validation dataset, which may be either genuinely new data, or a dataset obtained by splitting the original data into two parts (discovery and validation data) [11]. We use the latter approach in our study.

The connection between the multiplicity of analysis strategies and over-optimism is occasionally mentioned in the literature, mostly in relation to significance testing [12]. For example, it is well known that trying different analysis choices can make it easier to find a statistically significant result [7, 13]. If the researcher does this in an intentional manner (i.e., tweaking the analysis choices sequentially until a “significant” p-value is reached), this is called *p-hacking* [14]. However, over-optimistic bias might also appear without conscious “hacking”: A researcher may try different methods with the best intentions but then proceed to *selective reporting* (reporting only the method that yields the best result). Additionally, such effects do not only pertain to significance testing, but may appear whenever the result of a statistical analysis is quantified (e.g., with a performance measure or an index value).

In this paper we focus on over-optimism in the context of unsupervised microbiome analysis, outside of the classical setting of significance testing. We illustrate the over-optimistic effects caused by the multiplicity of analysis strategies in combination with selective reporting, as quantified by the subsequent validation of the optimistic results. As exemplary data, we use OTU count data from the American Gut Project (AGP) obtained with 16S amplicon sequencing [15]. It is well known that technical variation in amplicon sequencing (e.g., batch effects with respect to different labs or different machines) or using different methods for clustering sequences to obtain OTUs may lead to variation in the generation of the OTU count data and the results of subsequent statistical analysis [16, 17, 4, 18]. In the present work, however, we focus on the multiplicity of the statistical analysis methods (starting from the processed OTU count table), which has received somewhat less attention than multiplicity stemming from different technical methods. Recently, some studies have highlighted that different statistical analysis methods or modeling strategies may yield inconsistent results, namely in the context of microbiome-disease association modeling [19], microbiome differential abundance methods [20], and analyzing microbiome intervention design studies [21]. In contrast to these studies, a) we focus on the multiplicity of *unsupervised* statistical methods, i.e., methods for network learning and clustering, and b) our main goal is not to compare the results of different methods, but rather to quantify over-optimism effects that stem from picking the “best” result. The range of the statistical methods we consider includes 1) normalization to make read counts comparable across samples and to account for compositionality (if required by the subsequent analysis steps), 2) the generation of bacteria networks or distance matrices, and 3) methods to further process the network/distance information such as clustering.

The key idea of our illustrative study consists in splitting the whole dataset into a discovery and a validation set, trying out different methods for each of these three analysis steps on the discovery data, choosing the combination of methods that yields the best result on the discovery data according to an evaluation criterion, and applying this combination to the validation data to check whether the evaluation criterion takes a similar value. Fig 1 gives an overview of this approach, which we now describe in more detail.

**Fig 1.**
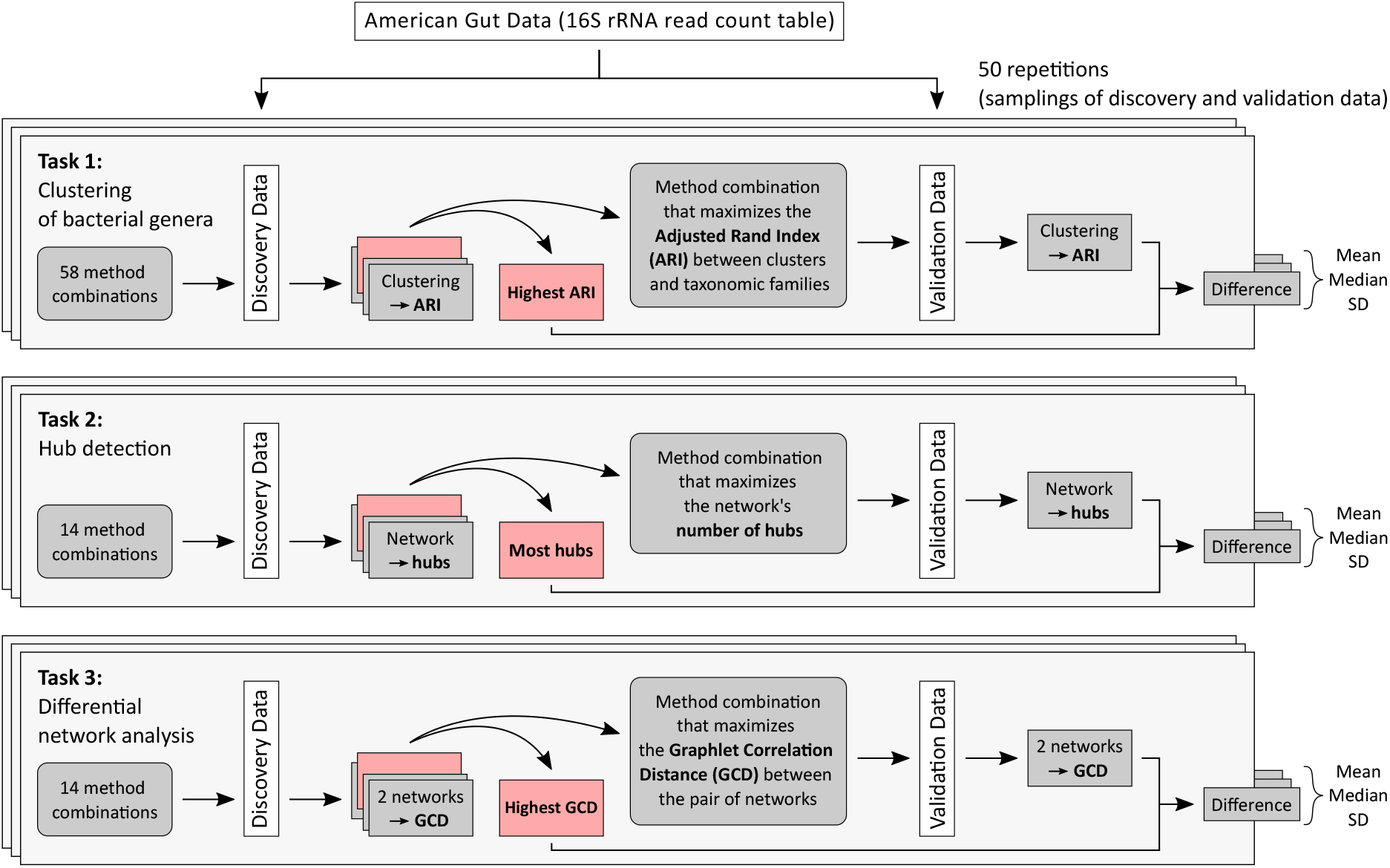
Graphical overview of our study. The process of drawing 50 samplings of discovery and validation data is repeated for different sample sizes: *n* ∈ {100, 250, 500, 1000, 4000} for the Tasks 1 and 2 and *n* ∈ {100, 250, 500} for Task 3.

We use three exemplary “research tasks” to illustrate the effects of the multiplicity of analysis strategies. Imagine a researcher who wishes to perform an unsupervised analysis of microbiome data. Even though the analysis is unsupervised, the researcher usually has some hopes for the results. While these expectations could be vague at first, the researcher might eventually focus on a concrete evaluation criterion that represents these hopes in order to judge the results. The researcher tries different statistical methods and chooses the method that yields the best result according to the evaluation criterion. We now detail the three research tasks, the hopes that our hypothetical researcher might have, and the concrete evaluation criteria they might use (and which we therefore choose for our illustrative study):

1. **Clustering of bacterial genera:** The hope is to find a clustering of bacterial genera that yields good agreement with the taxonomic categorization of the genera into families. As concrete evaluation criterion we choose the Adjusted Rand Index (ARI, [22]), a measure for comparing two partitions, normalized for chance agreement. The ARI ranges in [–1,1], with higher values indicating higher similarity of the partitions. In this research task, one partition is given by the clustering, the other one by the taxonomic categorization. The aim is not to find a clustering that is *perfectly* aligned with the taxonomic categorization – otherwise, the researcher could just use the categorization directly. Still, some agreement with the taxonomy is often considered as a good property of a clustering of bacteria [23].
2. **Hub detection:** A researcher might hope to find a microbial network with interesting keystone taxa (also called “microbial hubs”), i.e., highly connected taxa which are assumed to have a strong impact on the rest of the network. Detecting and analyzing keystone taxa in order to better understand microbial interactions has become popular in recent years [24, 25, 26]. In our illustrative example, we assume that our hypothetical researcher is interested in choosing a method that yields a relatively high number of hubs, to maximize the “hubbiness” of the network. Thus the number of hubs is used as the concrete evaluation criterion. Of course, other criteria to choose an “interesting” network with hubs are also feasible.
3. **Differential network analysis:** Microbiome researchers are often interested in the effects of treatments, such as antibiotics, on the gut microbial community (see, e.g., [27, 28] for background). When generating microbial association networks for two groups (one for persons who did not take antibiotics in the last year, and one for persons who took antibiotics in the last month), a researcher might expect that the networks (as proxies for microbial community structure) potentially change. As concrete evaluation criterion we measure the dissimilarity between the networks with the Graphlet Correlation Distance (GCD) [29]. The method that yields the largest GCD between the two networks is chosen. The GCD has been used in previous studies to compare microbial networks [30, 31, 32].

For each of the three research tasks, we emulate our “hypothetical researcher” by trying different methods (i.e., methods for generating microbial networks and/or clustering) and looking for the best result. The hypothetical researcher might stop at this point, and only report the best result according to the respective criterion. In contrast, we are interested in whether the best result can be confirmed on *validation data*: The result obtained by the “best” method on the discovery data (i.e., the “best” ARI, number of hubs, or GCD, respectively) is compared with the result obtained by this method on the validation data. The discovery and validation datasets are obtained by randomly sampling two disjoint subsets from the full AGP dataset, a process which is repeated multiple times.

Note that our analysis serves only illustrative purposes to study over-optimism effects. It is not our aim to systematically evaluate or compare the chosen method combinations. Moreover, we do not claim that researchers typically apply multiple methods to a dataset as systematically as we do this here, nor that they “optimize” for the best method with malicious intentions. Nevertheless, during a longer research process, researchers will often try multiple methods on a dataset, and even if this happens with the best intentions, it might still cause over-optimism effects.

We present the results of our analysis in Section 2. Section 3 contains a discussion. In Section 4, we give a detailed overview of the exemplary dataset, our study design, and the different statistical methods that we applied to the discovery data.

## 2 Results

### 2.1 Quantifying over-optimism effects

For each research task, we drew discovery and validation sets (each with sample size n) of varying sizes: *n* ∈ {100, 250, 500,1000, 4000} for the first two research tasks and *n* ∈ {100, 250, 500} for the third research task. In the latter case, the maximal sample size is reduced due to the required information about antibiotics usage (more details are given in Section 4.2). For each *n*, the process of drawing discovery and validation sets is repeated 50 times. As sampling variability decreases with increasing *n*, the performances of a method on both discovery and validation data should become more and more similar. We thus expected over-optimistic effects to decrease with increasing *n*.

For each research task, we applied multiple method combinations to the discovery data, involving *normalization methods* (clr [33], mclr [34], and VST [35]), *association estimation* (Pearson correlation, Spearman correlation, latentcor [36], SPRING [34], and proportionality [37]), *sparsification* (*t*-test, threshold method, and neighborhood selection), and, for the first research task, *clustering* (hierarchical clustering, spectral clustering [38], fast greedy modularity optimization [39], Louvain method for community detection [40], and manta [41]). Detailed descriptions of the combinations are given in Section 4.3.

Supplementary figures show the results of applying the different method combinations to the discovery data for the varying sample sizes (see the Supplementary Material). Notably, there is some change in the selected “best” method combination with respect to sample size. In particular, the performance of the sparsification methods is dependent on the sample size. These results are discussed in detail in the Supplementary Material.

Our main interest lies in choosing the method combination that yields the maximum value of the evaluation criterion (ARI, number of hubs, and GCD) on the discovery data, applying it to the validation data, and checking whether the values of the evaluation criteria can be validated. Over-optimism is indicated if the value of the evaluation criterion is lower on the validation data compared to the result on the discovery data. Exemplary results for *n* = 250 are shown in Fig 2. The corresponding figures for all other sample sizes *n* are given in the Supplementary Material.

**Fig 2.**
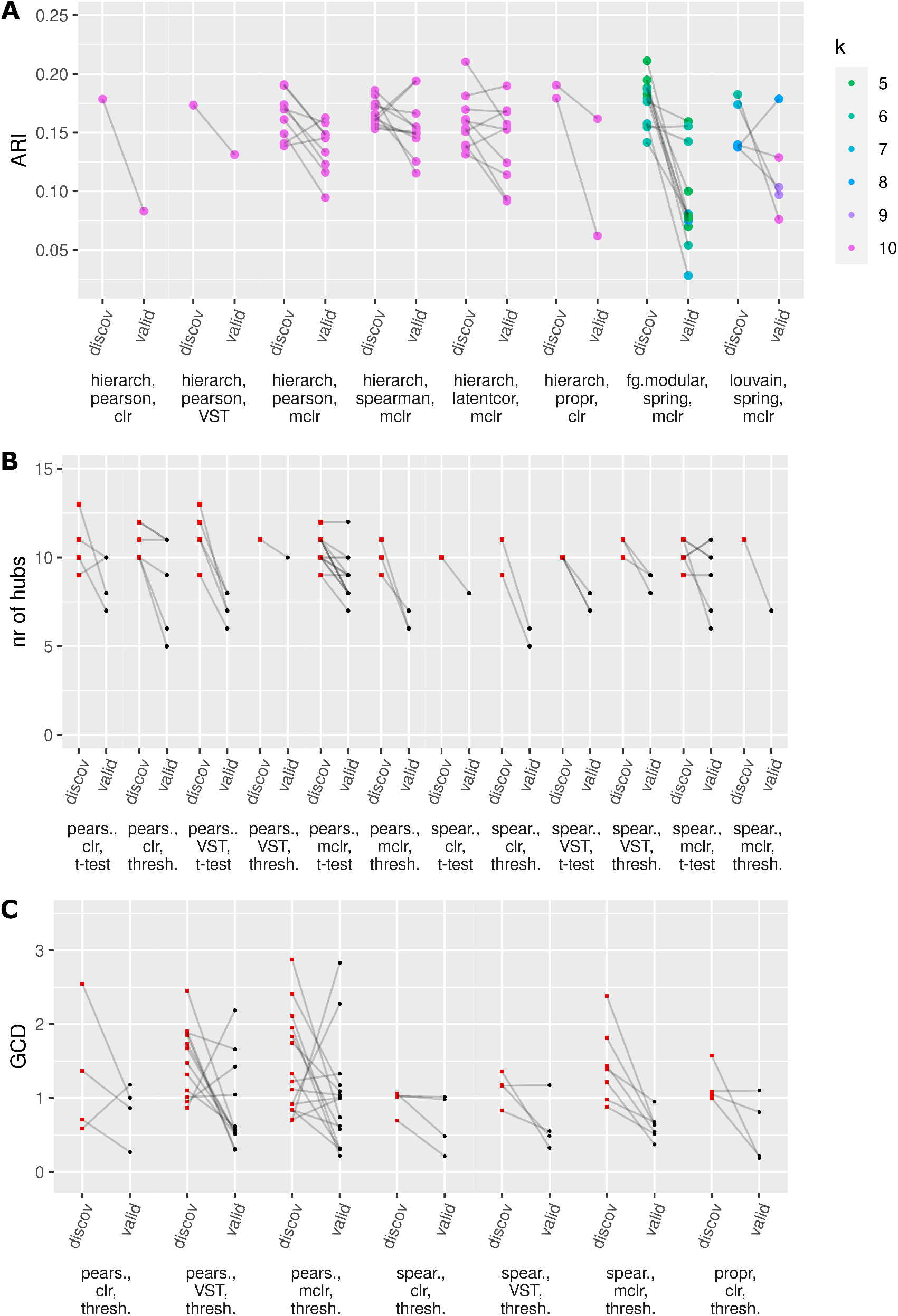
For *n* = 250: Values of the evaluation criteria resulting from the “best” method combinations on the discovery data, compared to the corresponding results on the validation data. a) ARI values for the clustering task, b) numbers of hubs for the hub detection task, c) GCD values for the differential network analysis task.

On the *x*-axis, only the method combinations that performed best in at least one of the 50 samplings are shown (that is, not *all* tried method combinations). For each sampling, the value of the evaluation criterion on the discovery data (belonging to the best method combination) and the corresponding value on the validation data are connected by lines. For the clustering task, the dots representing the ARI values are colored according to the number *k* of clusters in the respective clustering result. (For hierarchical and spectral clustering, we fixed the number of clusters at *k* = 10. The other clustering algorithms all have an inbuilt mechanism for determining *k*. Details are given in Section 4.3.1.) For the other two research tasks, the results are shown as red squares for the discovery data and black dots for the validation data.

The lines point downwards in most cases, i.e., the results for the validation data are usually slightly worse than for the discovery data. This indicates over-optimism effects. To further quantify these effects, Table 1 shows the mean, median, and standard deviation of the difference as well as the scaled difference between the value of the evaluation criterion on the validation data and the value on the discovery data (over the 50 samplings of discovery/validation data).

**Table 1.** Mean, median, and standard deviation (over 50 samplings of discovery/validation data) of the difference (both unscaled and scaled) between the value of the evaluation criterion on the validation data and the corresponding value on the discovery data. *ARI_discov_* denotes the best ARI on the discovery data and *ARI_valid_* the ARI resulting from the corresponding method combination on the validation data. The quantities *#hub_sdiscov_, #hubs_valid_* (number of hubs) and *GCD_discov_, GCD_valid_* are defined analogously.

| Research task 1: clustering of bacterial genera |  |  |  |  |  |  |
| --- | --- | --- | --- | --- | --- | --- |
| $n$ | $ARI_{valid} - ARI_{discov}$ | | | $\frac{ARI_{valid} - ARI_{discov}}{ARI_{discov}}$ | | |
|  | mean | median | sd | mean | median | sd |
| 100 | -0.054 | -0.046 | 0.044 | -30.0% | -26.7% | 24.1% |
| 250 | -0.039 | -0.035 | 0.046 | -22.0% | -21.8% | 26.3% |
| 500 | -0.038 | -0.037 | 0.038 | -21.4% | -20.6% | 21.6% |
| 1000 | -0.042 | -0.035 | 0.037 | -23.7% | -20.1% | 20.3% |
| 4000 | -0.035 | -0.033 | 0.035 | -19.0% | -18.3% | 18.8% |

| Research task 2: hub detection |  |  |  |  |  |  |
| --- | --- | --- | --- | --- | --- | --- |
| $n$ | $\#hubs_{valid} - \#hubs_{discov}$ | | | $\frac{\#hubs_{valid} - \#hubs_{discov}}{\#hubs_{discov}}$ | | |
|  | mean | median | sd | mean | median | sd |
| 100 | -2.44 | -3 | 2.35 | -21.6% | -24.0% | 21.3% |
| 250 | -2.18 | -2 | 1.78 | -20.5% | -20.0% | 16.5% |
| 500 | -2.12 | -2 | 1.88 | -20.8% | -20.0% | 17.9% |
| 1000 | -1.64 | -2 | 1.52 | -16.3% | -18.2% | 15.4% |
| 4000 | -1.12 | -1 | 1.32 | -11.5% | -11.1% | 13.8% |

**Table 1.** Mean, median, and standard deviation (over 50 samplings of discovery/validation data) of the difference (both unscaled and scaled) between the value of the evaluation criterion on the validation data and the corresponding value on the discovery data.
| Research task 3: differential network analysis |  |  |  |  |  |  |
| --- | --- | --- | --- | --- | --- | --- |
| $n$ | $GCD_{valid} - GCD_{discov}$ | | | $\frac{GCD_{valid} - GCD_{discov}}{GCD_{discov}}$ | | |
|  | mean | median | sd | mean | median | sd |
| 100 | -0.481 | -0.463 | 0.829 | -25.0% | -30.0% | 55.5% |
| 250 | -0.555 | -0.516 | 0.856 | -26.7% | -52.8% | 72.5% |
| 500 | -0.305 | -0.417 | 0.605 | -18.6% | -45.1% | 63.5% |

As expected, the means and medians of the differences are negative for all three research tasks and all sample sizes, demonstrating that the results on the discovery data were somewhat over-optimistic. We now discuss the behavior of the average differences over the varying sample sizes *n* in more detail for each research task in turn.

#### Research task 1 (clustering)

The average absolute decline of the ARI on the validation data is not drastic, but when considering the scaled difference, the ARI is reduced on the validation data by about 20-30% on average. Note that the absolute value of the mean/median ARI difference (both unscaled and scaled) is largest for *n* = 100, and smallest for *n* = 4000. This fits with our previously mentioned hypothesis that over-optimism effects are less pronounced when *n* is large. However, between 100 and 4000, there is no clear linearly decreasing tendency in the absolute mean/median ARI differences.

#### Research task 2 (hub detection)

The absolute values of the means and medians of the differences tend to decrease with increasing sample size. Again, this fits with our hypothesis that the over-optimistic bias decreases with increasing *n*.

#### Research task 3 (differential network analysis)

The absolute values of the means and medians do not monotonically decrease with increasing *n*: for *n* = 250, these are slightly larger than for *n* = 100. This is perhaps due to the fact that the sampling variability is still rather large at *n* = 250. At *n* = 500, however, the over-optimism effect appears to decrease, as evidenced by the drops in the absolute values of the average differences (both unscaled and scaled). For even higher sample sizes, we would expect to see a continuing decline of the over-optimistic bias, although we cannot confirm this due to the limited data availability.

### 2.2 Additional stability analyses

While our main focus was to compare the “best” result on the discovery data to the corresponding result on the validation data with respect to the evaluation criteria, we also compare these results with some additional stability evaluations (for the first two research tasks) to further demonstrate that the methods do not necessarily yield stable results on discovery vs. validation data. For the clustering task, we compared the clusterings on discovery vs. validation data with the ARI (while the agreement with the taxonomic categorization was ignored). This measure is denoted as *ARI_stab_*. The results are reported in Table 2. For the hub detection task, we compared the sets of hubs on discovery vs. validation data with the Jaccard index (on the genus level) and cosine similarity index (on the family level), as reported in Table 3. The indices are described in more detail in Section 4.3.

**Table 2.** Mean, median, and standard deviation of *ARI_stab_*, i.e., the ARI between the clusterings on discovery and validation data, over 50 samplings of discovery/validation data.

| $n$ | $ARI_{stab}$ | | |
| --- | --- | --- | --- |
|  | mean | median | sd |
| 100 | 0.361 | 0.329 | 0.111 |
| 250 | 0.509 | 0.491 | 0.166 |
| 500 | 0.604 | 0.574 | 0.168 |
| 1000 | 0.600 | 0.568 | 0.166 |
| 4000 | 0.763 | 0.792 | 0.140 |

**Table 3.** Mean, median, and standard deviation (over 50 samplings of discovery/validation data) of a) the Jaccard index which compares the set of hubs obtained on the discovery data with the set of hubs on the validation data, and b) the cosine similarity which compares these sets of hubs, but on the level of families.

| $n$ | Jaccard | | | Cosine similarity | | |
| --- | --- | --- | --- | --- | --- | --- |
|  | mean | median | sd | mean | median | sd |
| 100 | 0.236 | 0.250 | 0.109 | 0.881 | 0.911 | 0.112 |
| 250 | 0.359 | 0.357 | 0.119 | 0.922 | 0.955 | 0.078 |
| 500 | 0.443 | 0.429 | 0.116 | 0.948 | 0.969 | 0.060 |
| 1000 | 0.546 | 0.538 | 0.139 | 0.946 | 0.974 | 0.068 |
| 4000 | 0.709 | 0.727 | 0.147 | 0.975 | 0.984 | 0.026 |

For the clustering task, Table 2 shows that for smaller sample sizes, the mean ARIs are rather far away from 1, which indicates notable differences between the clusterings of the bacteria based on discovery vs. validation data. The clusterings tend to become more similar with increasing sample size, but even for *n* = 4000, the mean ARI of about 0.8 indicates that the clusterings are still different to some extent. This shows that the chosen clustering on the discovery data is not necessarily stable regarding cluster memberships when the result is validated on the validation data.

For the hub detection task, Table 3 demonstrates that the sets of hubs can be quite different between discovery and validation data, as measured with the Jaccard index (which ranges between 0 and 1). For smaller sample sizes, the similarity is particularly small. The Jaccard values increase with increasing sample size, but even at *n* = 4000, a mean value of about 0.7 shows that there are still notable dissimilarities between the sets of hubs. For the similarity on *family* level, we expect higher values (given that two hubs from the same family which differ on the genus level are counted as not equal for the Jaccard index and as equal for the cosine similarity). Indeed, the values of the cosine similarity (which ranges between −1 and 1), are generally quite high. Therefore, if one only interprets the hubs on family level (e.g., with respect to typical functions of the bacterial families), there is less danger of instability between discovery and validation data, compared to an interpretation on genus level.

## 3 Discussion

We have quantified over-optimism effects resulting from the multiplicity of analysis strategies coupled with selective reporting, using three exemplary microbiome research questions. Our results indicate an over-optimistic bias for all three research tasks. That is, when choosing the “best” method on the discovery data according to the maximization of an evaluation criterion, this criterion then tends to attain lower (“worse”) values on the validation data when the same method is applied. The exact size of the over-optimistic bias depends on the research task and sample size. Generally speaking, the over-optimistic bias tends to be more pronounced at smaller sample sizes, although the relation between sample size and optimistic bias is not always strictly monotonically decreasing in our analyses. Additional stability analyses for the first two research tasks have illustrated that clustering solutions and sets of hubs – which have been yielded by a method on discovery data – do not necessarily remain stable when the same method is applied to validation data.

In summary, our study has demonstrated that the issue of over-optimism and instability of results goes beyond the context of statistical testing and fishing for significance, and pertains to unsupervised analysis strategies as well. Over-optimism can lead to unreliable results, and might ultimately hinder research progress. We now discuss some strategies which may help researchers avoid over-optimistic bias in their application studies.

The first option is to reduce the multiplicity of analysis strategies *before* the start of the analysis. Researchers should carefully consider which method is most suitable for their application. Here, guidance from *neutral comparison studies* can be relevant. Such studies compare existing methods (instead of introducing a novel method), and the authors of the study are neutral, i.e., they do not have a vested interest in a particular method showing better performance than the others and are as a group approximately equally familiar with all considered methods. We refer to [42, 43] for a more detailed discussion of this concept. It would be desirable if more neutral comparison studies were published in the context of methodological research on microbiome analysis. For example, two recent studies already provide such a welcome effort in the context of microbial differential abundance testing [44, 20], and guidelines for benchmarking microbiome analysis methods have been proposed as well [45].

An additional strategy is *preregistration* of the researchers’ analysis plan. Preregistering refers to defining the research hypotheses and analysis plan, and posting this plan to a registry, *before* observing the results. This concept has gained plenty of attention in recent years [46]. Once their analysis plan is registered, researchers might shy away from trying many other analysis strategies and selectively reporting only the best results.

However, preregistration might not always be possible or sensible: for example, in exploratory research, researchers typically cannot pin down the exact analysis strategy in advance, and trying out different methods sequentially is quite natural [4]. Indeed, unsupervised analysis methods, on which we have focused in our study, are often used for exploratory purposes. In such cases, when the multiplicity of analysis strategies cannot be avoided, researchers should honestly report that their study is exploratory and that multiple methods were tried. They should not present their analyses as if a single analysis pipeline was fixed in advance, nor should they report only the “best” results.

In general, we would advise researchers to use validation data to validate their results whenever possible. While we have included validation data in our study to quantify over-optimism effects, researchers can also use validation data in their applied research, to check whether the best results on the discovery data still hold on the validation data. This is particularly relevant when the multiplicity of possible analysis strategies cannot be reduced beforehand, e.g., in the absence of relevant neutral comparison studies for the methods of interest. For the topic of cluster analysis (research task 1), different strategies for validating clustering results on validation data have been previously discussed in detail [11]. More awareness for the importance of validation data has also emerged in microbiome research (see, e.g., in the context of supervised analysis [47, 48] and large-scale cohort studies [49]).

In summary, we hope that our study helps raise awareness of the important problem of over-optimism in microbiome research, and that it motivates more widespread implementation of strategies to avoid over-optimistic bias. If researchers adhere to good research practices, the results of microbiome analyses will hopefully become more reliable and replicable in the future.

## 4 Materials and Methods

### 4.1 Dataset

We used data from the American Gut Project [15], a large citizen-science initiative. The project collected (mainly) fecal samples from participants in the United States, United Kingdom, and Australia. The researchers also collected metadata on the participants, e.g., health status, disease history, and lifestyle variables. Bacterial abundances were obtained using high-throughput amplicon sequencing, targeting the V4 region of the 16S rRNA marker gene with subsequent variant calling.

We downloaded an OTU count table for unrarefied bacterial fecal samples (dating from 2017) from the project website ftp://ftp.microbio.me/AmericanGut/ag-2017-12-04/, together with metadata about the samples. The OTU count table originally contained *p* = 35511 OTUs and *N* = 15148 samples. Following [23], we performed three preprocessing steps: 1) removing samples with a sequencing depth of less than 10000 counts, 2) removing OTUs which were present in less than 30% of the remaining samples, 3) removing 10% of the remaining samples, namely the samples with a sequencing depth under the 10%-percentile. The resulting OTU count table comprises *p* = 531 OTUs and *N* = 9631 samples.

For all three research tasks, the analysis is performed on the taxonomic rank of genera, to which the data were agglomerated. OTUs with unknown genus were assigned their own individual genus, which resulted in *p* = 323 genera overall.

### 4.2 Sampling of discovery and validation datasets

We obtained discovery and validation datasets by randomly sampling two disjoint subsets from the full AGP dataset. For each research task, this sampling process was performed along the *samples* of the AGP data (i.e., the subjects), not along the bacteria. This is because in each task, the bacteria formed a fixed set of entities of specific interest. This set thus remained constant for both discovery and validation data. For clustering, this is discussed in more detail in [11].

Discovery and validation sets (each with sample size n) were drawn of varying sizes: *n* ∈ {100, 250, 500, 1000, 4000} for the first two research tasks (clustering and hub detection), and *n* ∈ {100, 250, 500} for the third research task (differential network analysis). In the latter case, the maximal sample size was reduced because we only considered samples which did not take antibiotics in the last year as well as samples which took antibiotics in the last month. There were 6901 samples which fulfilled these criteria. Moreover, the sampling was stratified according to antibiotics use; for discovery and validation data each, we drew *n*/2 samples which did not take antibiotics in the last year and *n*/2 samples which took antibiotics in the last month. Because there are only 544 persons who took antibiotics in the last month, the maximum *n* is reduced to 500.

### 4.3 Methods for unsupervised microbiome analysis

In this section, we discuss which method combinations were applied to the discovery data, and how the results were evaluated on the validation data.

#### 4.3.1 Research task 1: Clustering bacterial genera

We varied different steps of the cluster analysis process, resulting in 58 method combinations that were tried on the discovery data. In this section we explain how the 58 combinations were obtained.

We used cluster algorithms from two categories. Algorithms from the first category are based on (dis)similarity matrices: hierarchical clustering and spectral clustering [38]. Algorithms from the second category are based on networks with weighted edges: fast greedy modularity optimization [39], the Louvain method for community detection [40], and the manta algorithm [41].

To generate either (dis)similarity matrices or weighted networks, associations (*r_ij_*)_*i,j*_ between the microbes must be calculated. Beforehand, often zero handling and normalization of the data are required. Table 4 gives an overview of the method combinations used for calculating the associations *r_ij_* for later use in (dis)similarity based clustering, i.e., for generating (dis)similarity matrices which will later be used as input for hierarchical and spectral clustering. We used four different association measures. The first ones are the Pearson and Spearman correlations, which require normalization to account for compositionality. Here we used either the centered log-ratio transformation (clr, [33]), the modified clr transformation (mclr, [34]), or the variance-stabilizing transformation (VST, [35]). As the clr and VST methods cannot handle zeros in the count data, a pseudo count of 1 was added to the count data before normalizing with these methods (mclr, on the other hand, can deal with zeros). Apart from the Pearson and Spearman correlations, we used the semi-parametric rank-based correlation, which is based on estimating the latent correlation matrix of a truncated Gaussian copula model (latentcor, [36, 50]). The latentcor method requires normalized counts that are strictly positive, and was therefore combined only with the mclr normalization. The final association measure is proportionality [37, 51]. While proportionality is a compositionally aware method, the data should still be log-ratio transformed before applying the proportionality measure, such that associations between different pairs of taxa are on the same scale and thus comparable [37]. We use the clr transformation as proposed in [37] and replace zero counts again by a pseudo count.

**Table 4.** Method combinations for generating microbial associations, which are then transformed into (dis)similarity matrices. The (dis)similarity matrices were used as input for hierarchical and spectral clustering.

| Zero handling | Normalization | Association estimation |
| --- | --- | --- |
| pseudo | clr | Pearson |
| pseudo | VST | Pearson |
| none | mclr | Pearson |
| pseudo | clr | Spearman |
| pseudo | VST | Spearman |
| none | mclr | Spearman |
| none | mclr | latentcor |
| pseudo | clr | proportionality |

Table 5 shows the method combinations used for calculating the associations *r_ij_* for later use in network-based clustering. That is, these methods were used for generating weighted networks. The method combinations are very similar to the methods in Table 4 for generating (dis)similarity matrices. Indeed, weighted networks are also based on (dis)similarity matrices, but the generation contains an additional sparsification step, as explained below. Again, the Pearson and Spearman correlations were used with the respective normalization and/or zero handling methods. We also used the SPRING method [34], which combines the latentcor correlation estimation with sparse graphical modeling techniques, namely by using the neighborhood selection technique [52] for sparse estimation of partial correlations. Finally, we used the proportionality measure.

**Table 5.** Method combinations for generating weighted microbial association networks. The networks were used as input for fast greedy modularity optimization, Louvain community detection, and manta.

| Zero handling | Normalization | Association estimation | Sparsification |
| --- | --- | --- | --- |
| pseudo | clr | Pearson | $t$-test |
| pseudo | clr | Pearson | threshold |
| pseudo | VST | Pearson | $t$-test |
| pseudo | VST | Pearson | threshold |
| none | mclr | Pearson | $t$-test |
| none | mclr | Pearson | threshold |
| pseudo | clr | Spearman | $t$-test |
| pseudo | clr | Spearman | threshold |
| pseudo | VST | Spearman | $t$-test |
| pseudo | VST | Spearman | threshold |
| none | mclr | Spearman | $t$-test |
| none | mclr | Spearman | threshold |
| none | mclr | SPRING | neighborhood selection |
| pseudo | clr | proportionality | threshold |

To generate a weighted network, the associations *r_ij_* (which are usually different from zero) were not directly used as an adjacency matrix – otherwise, the network would be dense. Therefore, the associations *r_ij_* were transformed into sparsified values 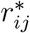 by setting some 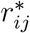 to zero to indicate that *i* and *j* are not connected, 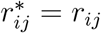 otherwise. For sparsification of the Pearson and Spearman correlations *r_ij_*, we used either Student’s *t*-test or the threshold method. The former sets 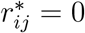 if the association *r_ij_* is not significantly different from 0 according to the *t*-test. The *p*-values were adjusted for multiple testing via the local false discovery rate [53]. For the threshold method, we set 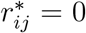 if *r_ij_* < *c* for some fixed threshold value *c* (we use *c* = 0.15 which gave reasonable results in preliminary analyses, not shown). For the proportionality measure, we used threshold sparsification. SPRING already comes with inbuilt sparsification given by the neighborhood selection method.

After calculating the associations as in Tables 4 and 5, they were then transformed as follows (the pipeline and notations are taken from [9]):

a. For (dis)similarity based clustering (Table 4): A dissimilarity matrix *D* = (*d_ij_*) for hierarchical clustering is calculated via 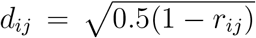. A similarity matrix *S* = (*s_ij_*) for spectral clustering is obtained by setting *s_ij_* = 1 – *d_ij_*.
b. For network-based clustering (Table 5): A weighted network is constructed as follows. For the edges *ij* with 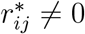 (i.e., the edges that remain after sparsification), the distances *d_ij_* and similarities *s_ij_* are calculated as in a). Finally, the weighted network is represented as adjacency matrix *A* = (*a_ij_*) with *a_ij_* = *s_ij_* for *ij* with 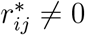, and *a_ij_* = 0 otherwise.

The (dis)similarity matrices and networks were then used as input for clustering. For hierarchical and spectral clustering, we fixed the number of clusters at *k* =10, which was inspired by the ten different taxonomic classes in the data. Also, *k* = 10 tends to yield better ARI results than *k*s lower than ten (preliminary analysis, not shown). *k*s higher than ten were not tried because we aimed to emulate a researcher who wants to find an interpretable, handy clustering (there are 34 different taxonomic families, but 34 clusters are not easily interpretable). The other clustering algorithms all have inbuilt mechanisms for determining *k*. Forcing *k* to be 10 for these methods generally did not improve the results (not shown). However, *k* can be indirectly influenced via the sparsification: The sparser the network, the more clusters tend to be found. This is one of the reasons we set the threshold for threshold sparsification at *c* = 0.15 because this value generally yielded sufficiently high *k*s to find good results, but only rarely *k*s that are so high that the clusters are difficult to interpret.

Overall, the method combinations yielded 58 different clustering results on the discovery data: 16 based on (dis)similarity clustering (8 rows in Table 4 times 2 cluster algorithms) and 42 based on network clustering (14 rows in Table 5 times 3 cluster algorithms). The best one out of the 58 clustering results is chosen, i.e., the clustering with the highest ARI regarding the taxonomic categorization into families. The corresponding method combination was applied to the validation data. The ARI between the clustering on the validation data and the taxonomic categorization was computed and compared with the best ARI on the discovery data. If the ARI on the validation data was lower, this was an indication that the best ARI on the discovery data was over-optimistic.

As an additional stability analysis, we compared the chosen clustering on the discovery data with the clustering on the validation data, again using the ARI.

#### 4.3.2 Research task 2: Hub detection

Here, we wanted to generate sparse weighted microbial association networks. For this purpose, we used the same methods as in Table 5. Thus, 14 method combinations were tried on the discovery data.

For hub detection in the resulting networks, hubs were defined as nodes that have the highest degree, betweenness, and closeness centrality [25]. More precisely, we determined the hubs as the nodes with centrality values above the 95% empirical quantile, for each of the three centrality measures simultaneously. The centralities are defined as follows [54]: The degree centrality denotes the number of adjacent nodes. The betweenness centrality measures the fraction of times a node lies on the shortest path between all other nodes. The closeness centrality of a node is the reciprocal of the sum of shortest paths between this node and all other nodes. All centrality measures were normalized to be comparable between networks of different sizes (see [9] for details). The centralities were only calculated for the largest connected component of each network (i.e., the largest subgraph of the network in which all nodes are connected); centrality values of nodes in the disconnected component were set to zero. We assumed that “hubs” in small parts of the network that are disconnected from the majority of the nodes are of less interest to researchers. Moreover, the betweenness and closeness centrality depend on shortest paths, which are not well-defined for nodes in different unconnected sub-graphs.

After applying the 14 method combinations and calculating the hubs for each resulting network, the method combination that yielded the highest number of hubs was chosen. If there were multiple method combinations that attained the maximal number of hubs, we chose the combination that yielded higher mean centrality values of the hubs. More specifically, for each set of hubs that corresponds to a method combination, the mean values of the three centrality measures were calculated over the hubs. Then for each centrality measure separately, the sets of hubs were ranked according to these mean values. Finally, the set of hubs (and thus the corresponding method combination) that yielded the highest mean rank over all three centrality measures was chosen.

The “best” method combination was then applied to the validation data. The number of hubs in the microbial network on the validation data was calculated and compared with the highest number of hubs on the discovery data. Over-optimism was indicated if the number of hubs was lower on the validation data.

Additionally, we reported the similarity of the sets of hubs determined on the discovery vs. validation data with the Jaccard index [55]: let *H_discov_, H_valid_* be the sets of hubs for the discovery resp. validation data, then

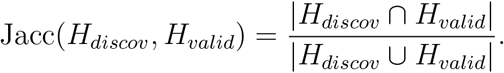

The Jaccard index takes values in [0,1], and is closer to 1 the more similar the sets are. The similarity between the sets of hubs was also assessed on the higher taxonomic level of families with the cosine similarity index. More precisely, assume that the hubs (genera) in the union *H_discov_* ∪ *H_valid_* belong to *l* distinct families overall. Let 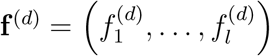 be the family frequency vector for *H_discov_*, that is, each entry 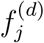 counts how many hubs in *H_discov_* belong to family *j*. Analogously, let **f**^(*v*)^ be the family frequency vector for *H_valid_*. The vectors **f**^(*d*)^ and **f**^(*v*)^ are then compared with the cosine similarity index:

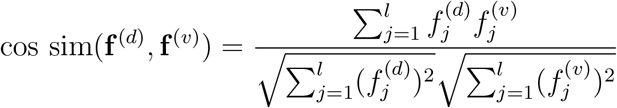

The cosine similarity index ranges in [0,1], with higher values indicating higher similarity.

#### 4.3.3 Research task 3: Differential network analysis

As described in Section 4.2, the discovery and validation datasets each consisted of two halves: persons who did not take antibiotics in the last year (“non-antibiotics samples”), and persons who took antibiotics in the last month (“antibiotics samples”). The methods for generating weighted microbial association networks as in Table 5 were applied separately to the antibiotics and non-antibiotics samples of the discovery data.

The resulting networks were compared with the Graphlet Correlation Distance (GCD, [29]). This distance measures the similarity of the networks based on small induced subgraphs, so-called graphlets. All graphlets composed of up to four nodes are considered, and the automorphism orbits of these graphlets are enumerated (orbits represent the “roles” that nodes can play in the graphlets). For each node in a given network, one can count how often the node participates in each graphlet at the respective orbits. Only 11 non-redundant orbits are considered here. Based on these orbit counts across all nodes, the 11 × 11 Spearman correlation matrix among the 11 orbits is calculated, which represents a robust and size independent network summary statistics. For comparing two networks, the Spearman correlation matrix is calculated for each network in turn. Then the Euclidean distance between the upper triangular parts of these matrices is calculated, resulting in the GCD.

In our study, the network generation method that yielded the largest GCD between the antibiotics network and the non-antibiotics network was chosen as the “best” one and applied to the antibiotics and non-antibiotics samples in the validation data. Again, the resulting networks are compared with the GCD. If the GCD on the validation data is smaller (i.e., the antibiotics vs. non-antibiotics networks are more similar than on the discovery data), this indicates over-optimism.

### 4.4 Technical implementation

All analyses were performed with R, version 4.0.4 and Python, version 3.6.13. Our fully reproducible code is available at https://github.com/thullmann/overoptimism-microbiome. (Dis)similarity matrices and weighted networks were generated with the R package Net-CoMi [9]. Spectral clustering was performed with a previously published R implementation [23]. For fast greedy modularity optimization and the Louvain method for community detection, we used the R package igraph [56]. For clustering with manta, we accessed the Python implementation [41] with the reticulate interface for R [57]. Orbit counts for the calculation of the GCD were generated with the R package orca [58].

## Supporting information

Supplementary results

## Acknowledgments

We thank Raphael Rehms for helpful comments about the Python code and Anna Jacob for valuable language corrections. This work has been partially supported by the German Federal Ministry of Education and Research (BMBF) [grant number 01IS18036A to Anne-Laure Boulesteix (Munich Center of Machine Learning)] and the German Research Foundation [grant number BO3139/7-1 to Anne-Laure Boulesteix]. The authors of this work take full responsibility for its content.

