## Supplementary results for "Over-optimism in unsupervised microbiome analysis: Insights from network learning and clustering"

### A Appendix: Supplementary results

#### A.1 Research task 1: Clustering of bacterial genera

Fig A.1-A.5 show the results of applying different clustering methods to the discovery data for sample sizes $n \in \{100, 250, 500, 1000, 4000\}$. For each method combination on the $x$-axis, the resulting ARIs, which measure agreement with the taxonomic categorization into families, are summarized over the 50 samplings by boxplots. Outliers are indicated by black crosses. Additionally, all results are shown as colored dots, with the color indicating the number $k$ of clusters in the respective clustering result. Results that were picked as the "best result" in one of the 50 samplings are marked by red square edges. For the network-based clustering methods with the networks generated by either the Pearson or Spearman correlation, the results for $t$-test and threshold sparsification are displayed together, i.e., 50*2 = 100 results are shown for these method combinations.

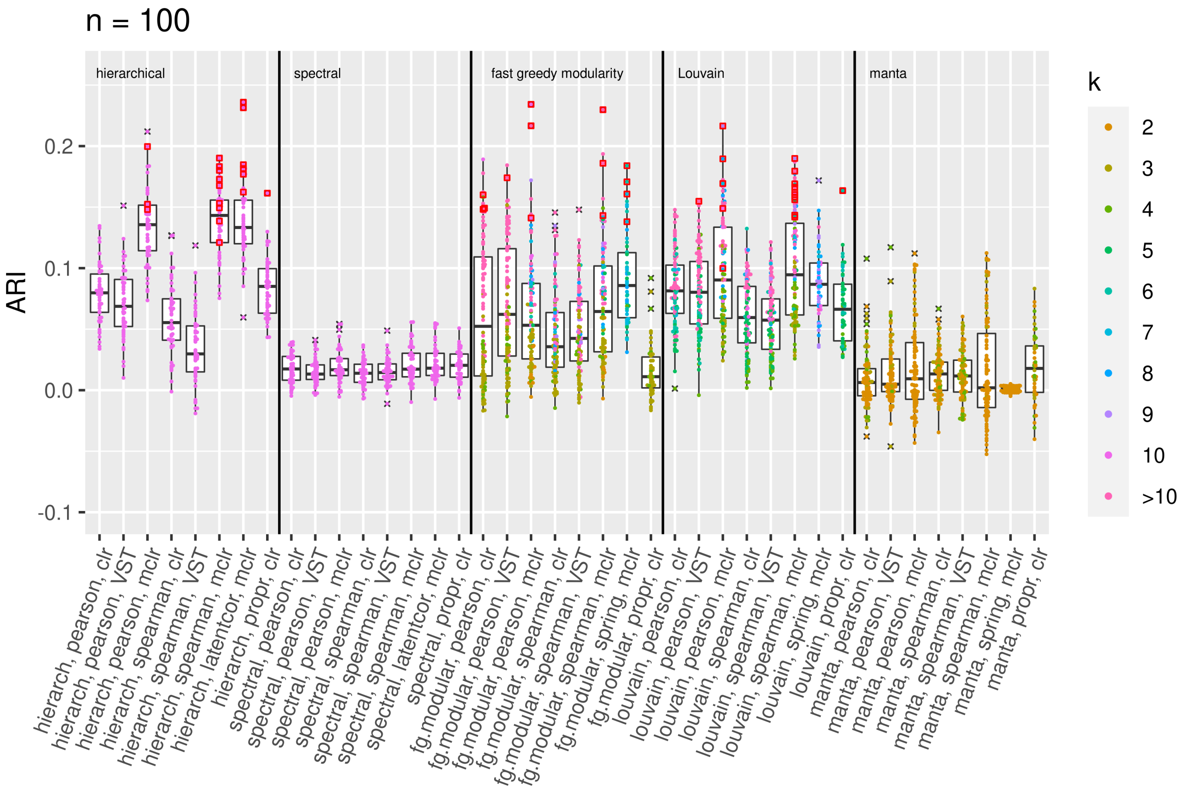


**Fig A.1.** Results for clustering bacterial genera on the discovery data, $n = 100$

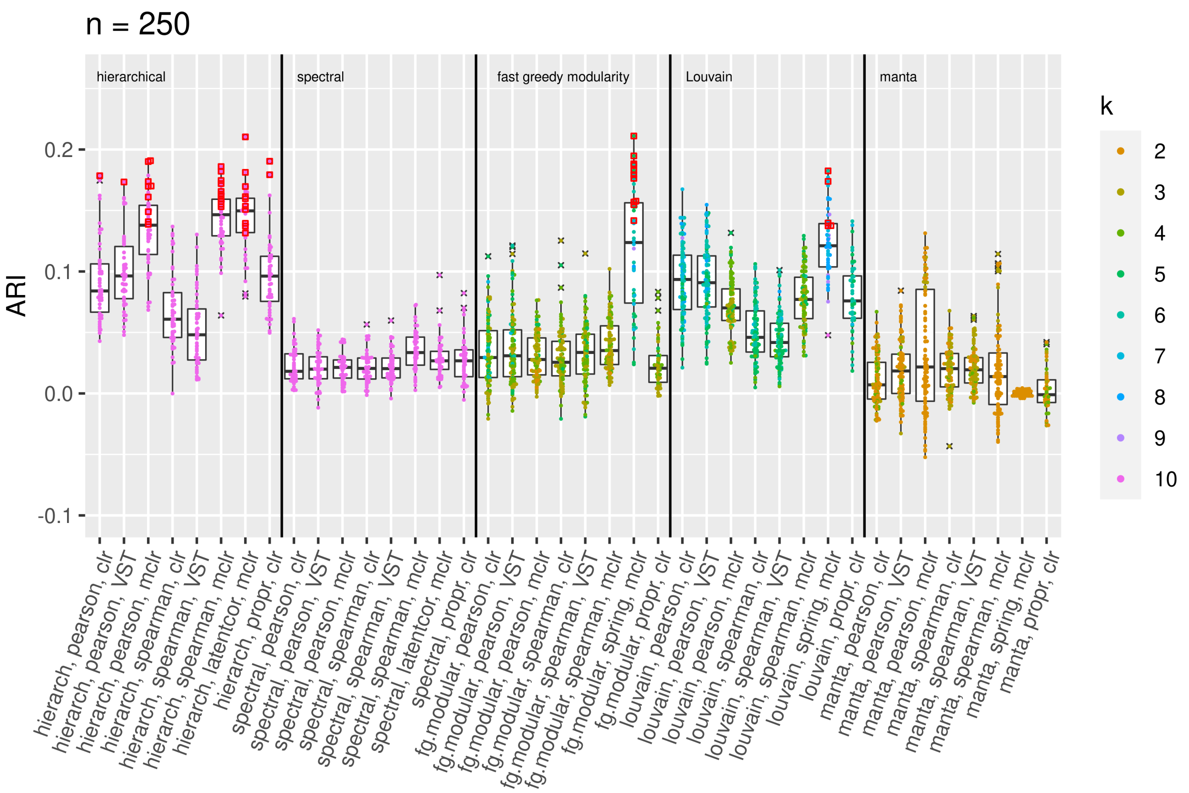


**Fig A.2.** Results for clustering bacterial genera on the discovery data, $n = 250$

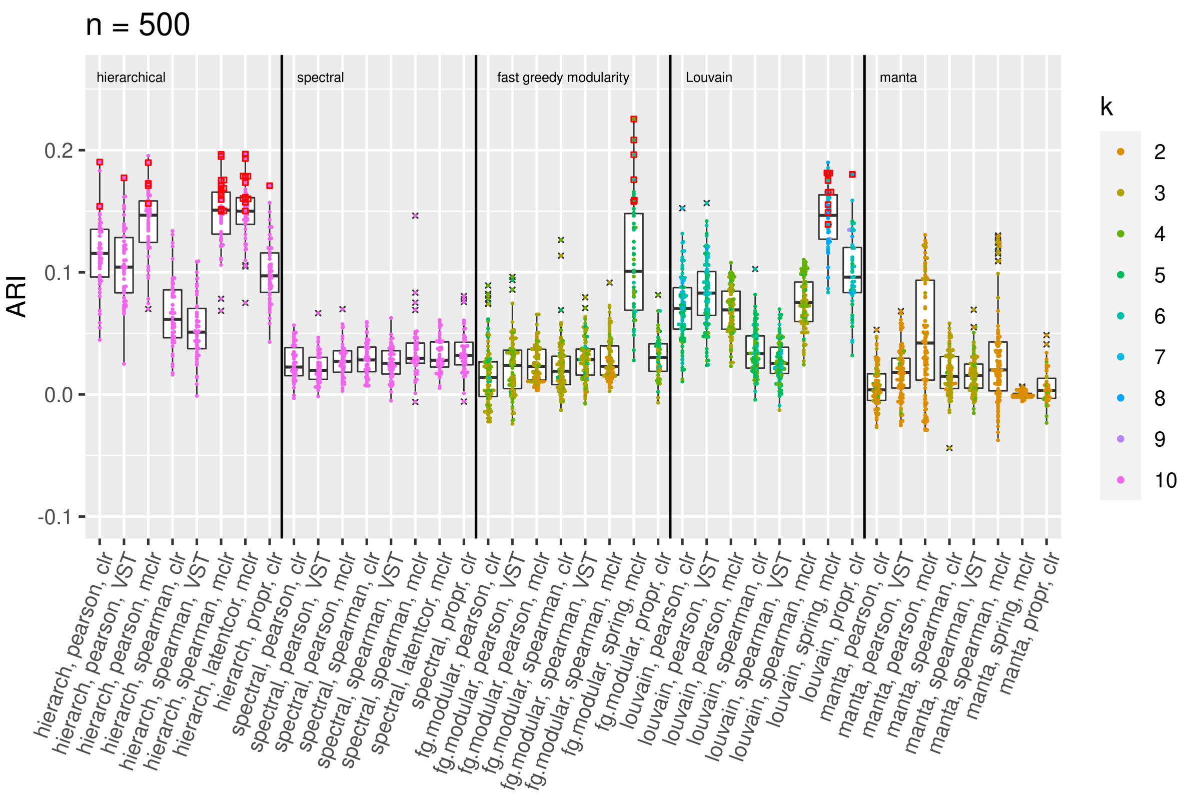


**Fig A.3.** Results for clustering bacterial genera on the discovery data, $n = 500$

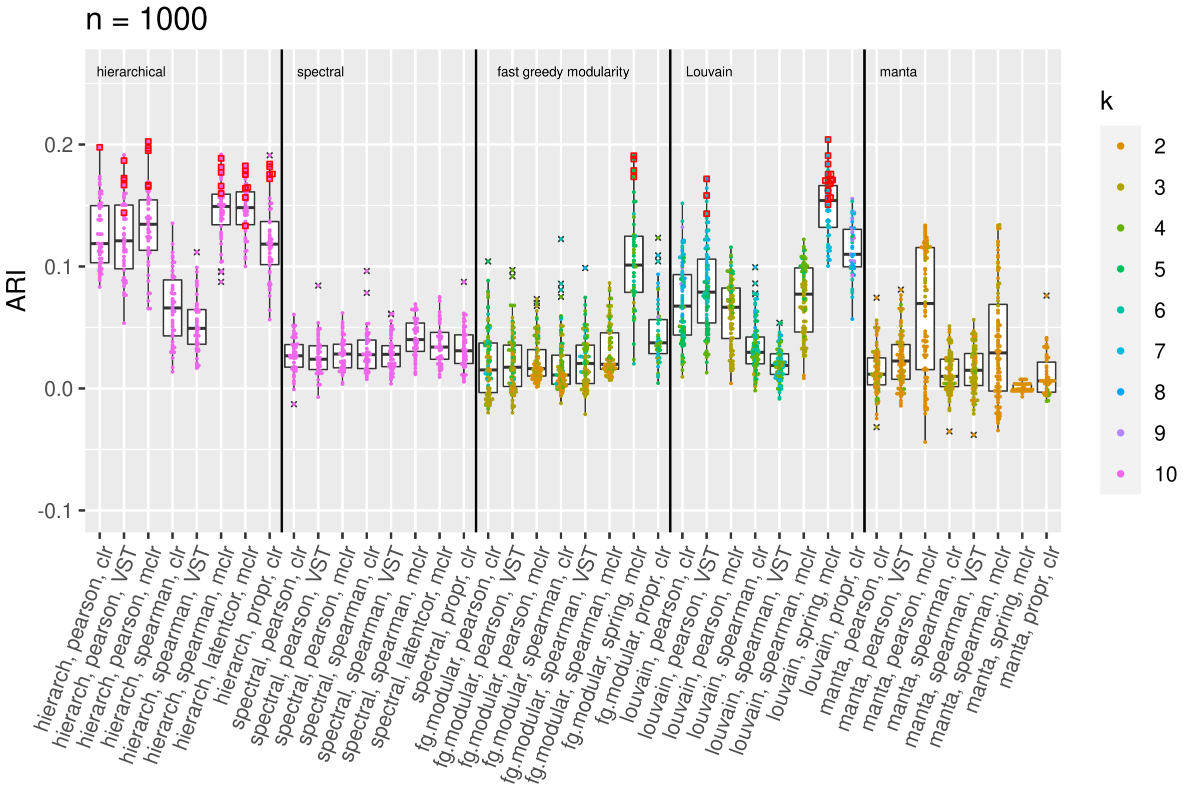


**Fig A.4.** Results for clustering bacterial genera on the discovery data, $n = 1000$

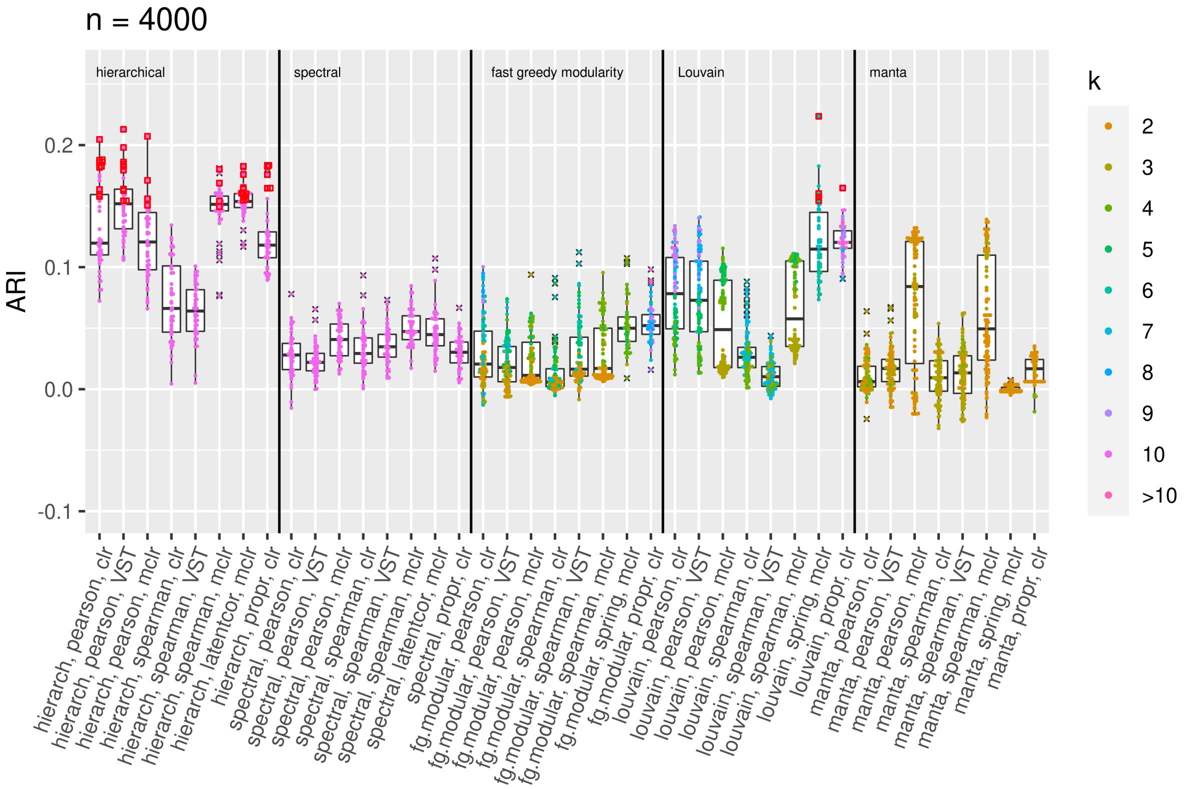


**Fig A.5.** Results for clustering bacterial genera on the discovery data, $n = 4000$

As can be seen in Fig A.1-A.5, the best ARI results stem either from hierarchical clustering, the Louvain method or fast greedy modularity optimization. Spectral clustering and manta are never selected. There is some change in the selected "best" methods with respect to sample size. For example, for $n = 100$, fast greedy modularity clustering performs well in several of the 50 samplings, but this cluster method does not yield very good ARI results for $n = 4000$. At $n = 4000$, hierarchical clustering is chosen as the best method in 45 of the 50 samplings.

In Fig A.6-A.10, results for the network-based clustering are shown separately for both sparsification methods ($t$-test and threshold). Results that were picked as the "best result" in one of the 50 samplings are marked by red square edges.

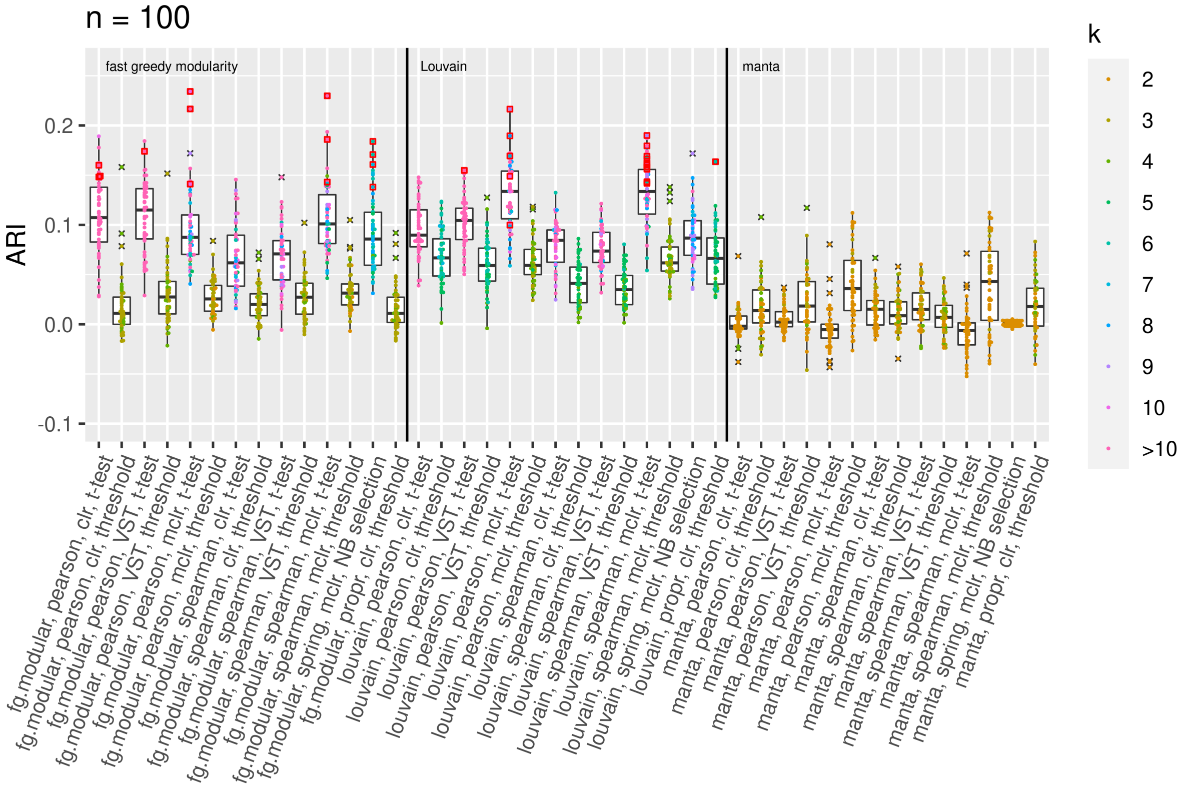


**Fig A.6.** Results for network-based clustering of bacterial genera on the discovery data, separated by sparsification methods, $n = 100$

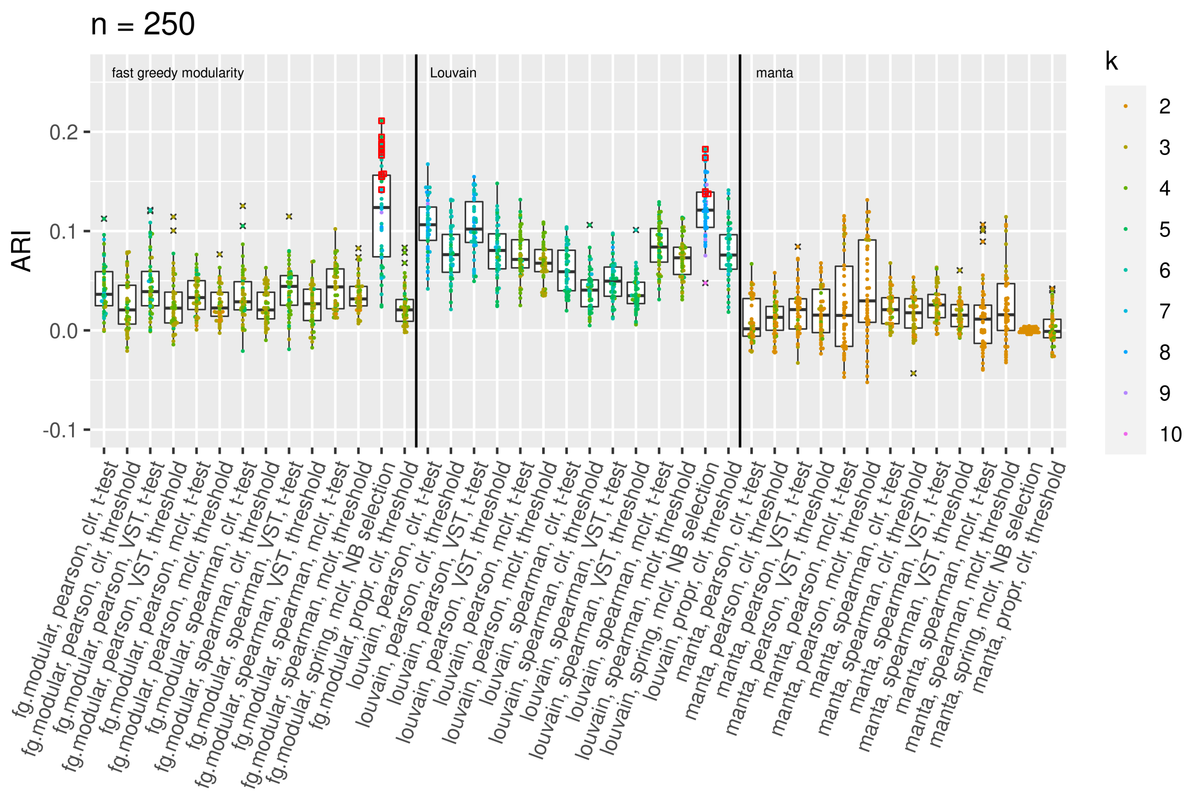


**Fig A.7.** Results for network-based clustering of bacterial genera on the discovery data, separated by sparsification methods, $n = 250$

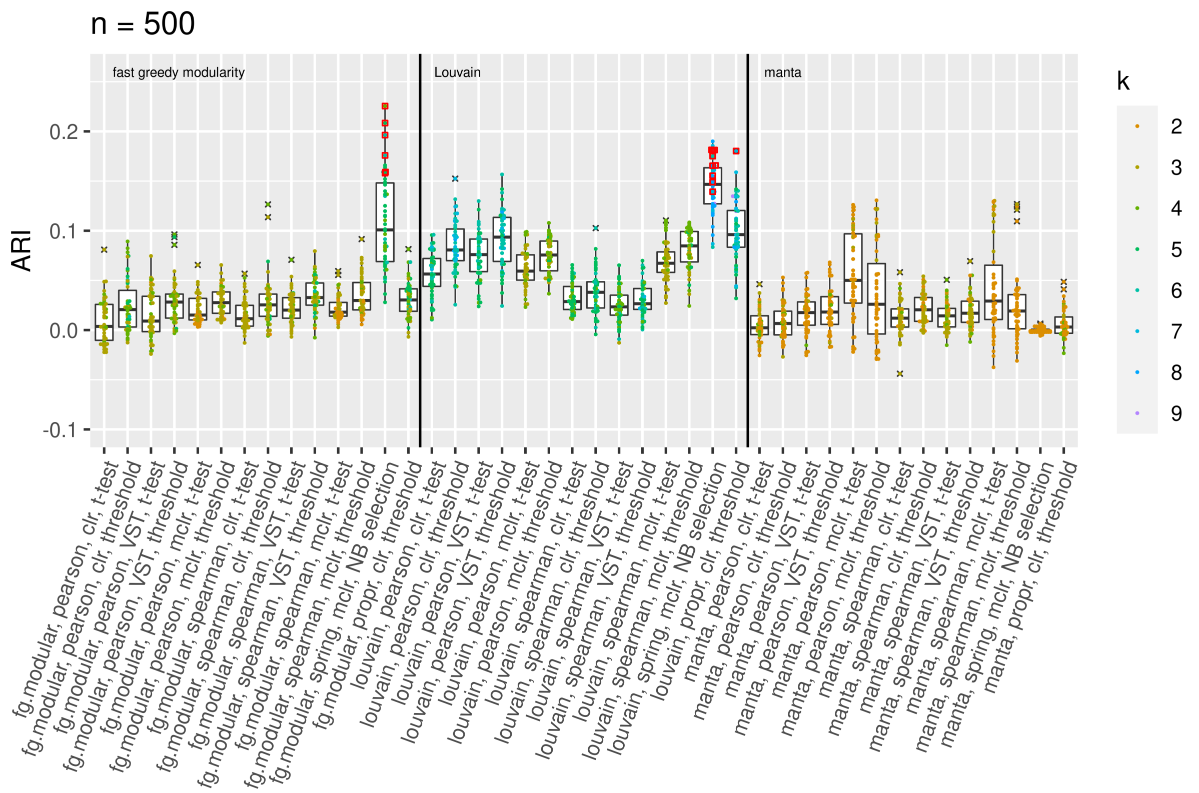


**Fig A.8.** Results for network-based clustering of bacterial genera on the discovery data, separated by sparsification methods, $n = 500$

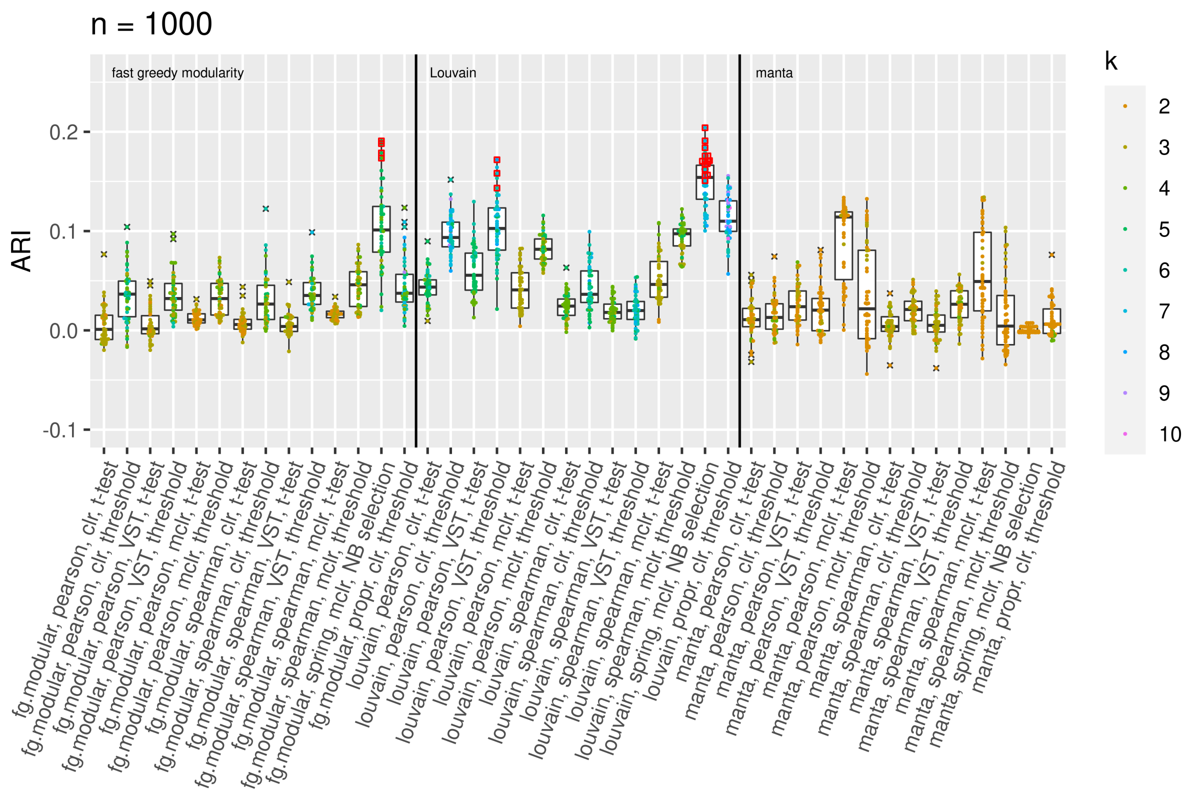


**Fig A.9.** Results for network-based clustering of bacterial genera on the discovery data, separated by sparsification methods, $n = 1000$

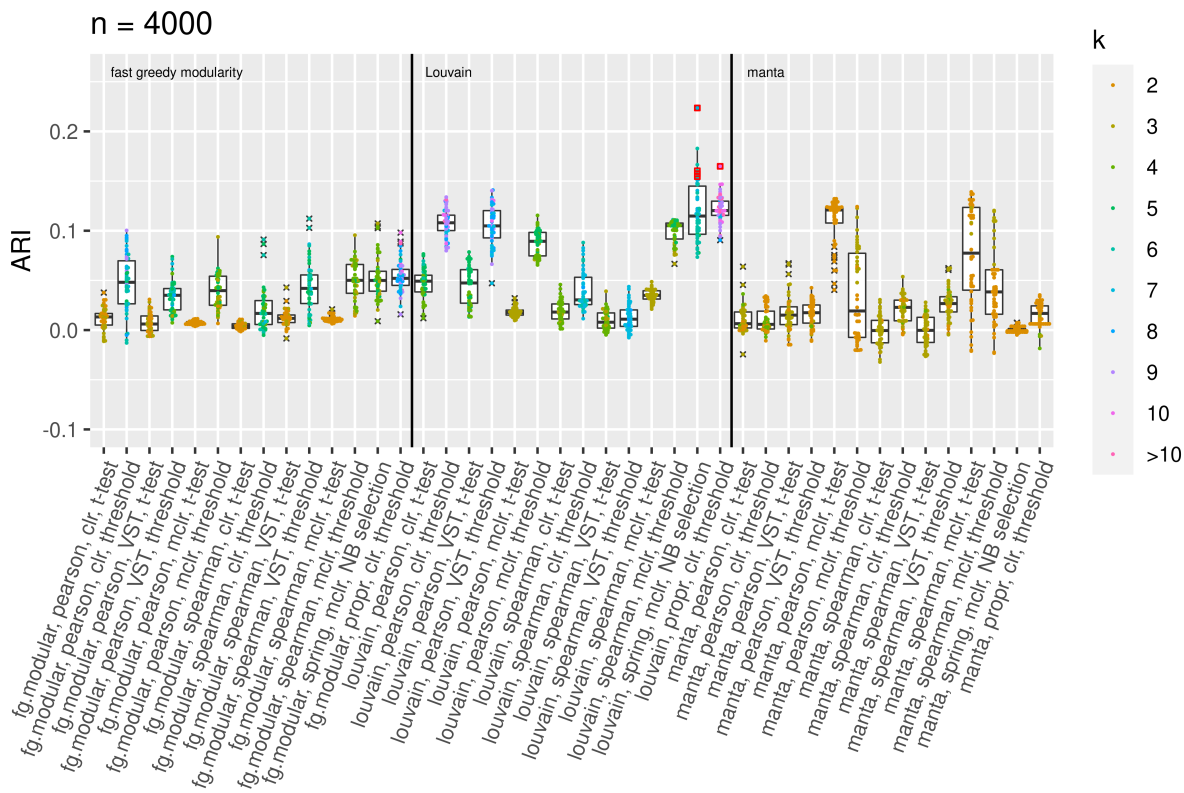


**Fig A.10.** Results for network-based clustering of bacterial genera on the discovery data, separated by sparsification methods, $n = 4000$

Our main interest lies in applying the "best" method to the validation data and checking whether the ARI result can be validated. The results are shown in Fig A.11-A.15. On the $x$-axis, the method combinations that were best in at least one of the 50 samplings are shown. The ARI values are shown as colored dots, with the color indicating the number $k$ of clusters in the respective clustering result.

For each of the 50 samplings, the respective best method combination is applied to the validation data. The ARI value on the discovery data (belonging to the best method combination) and the corresponding ARI on the validation data are connected by lines. The lines point downwards in most cases, i.e., the results for the validation data are usually slightly worse than for the discovery data.

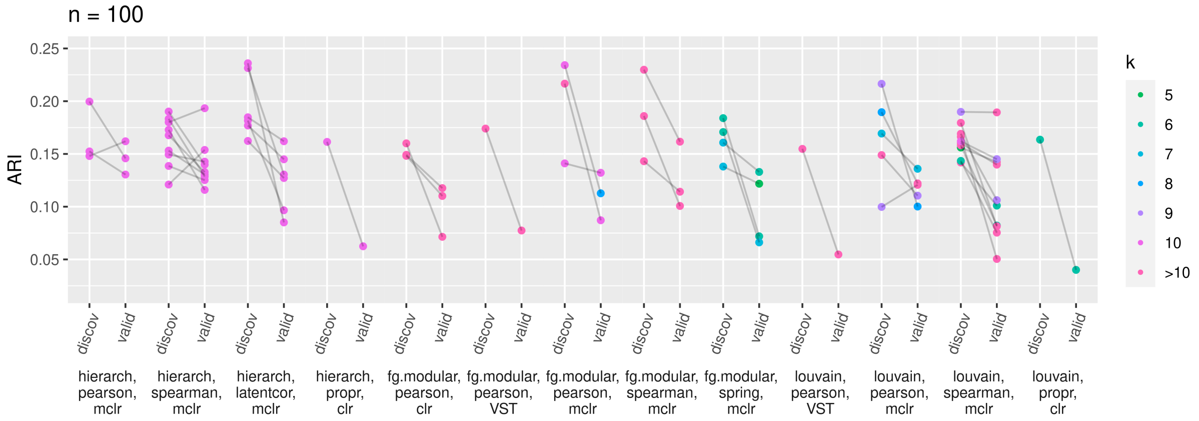


**Fig A.11.** Best ARIs for the clustering of bacterial genera on the discovery data, compared with the results on validation data, $n = 100$

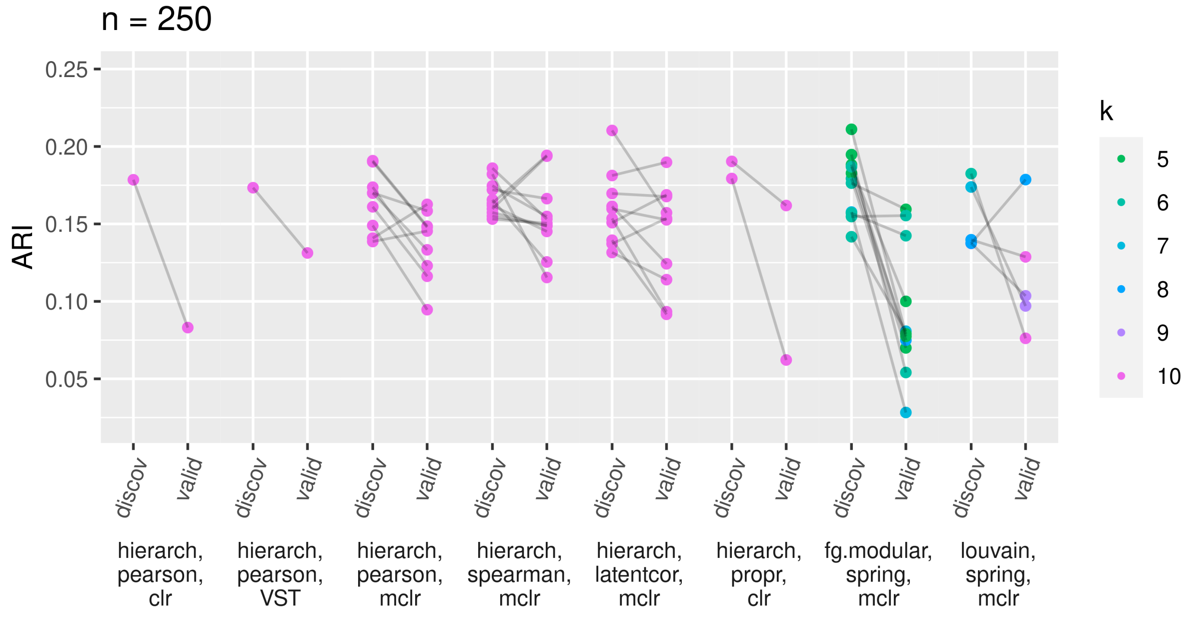


**Fig A.12.** Best ARIs for the clustering of bacterial genera on the discovery data, compared with the results on validation data, $n = 250$

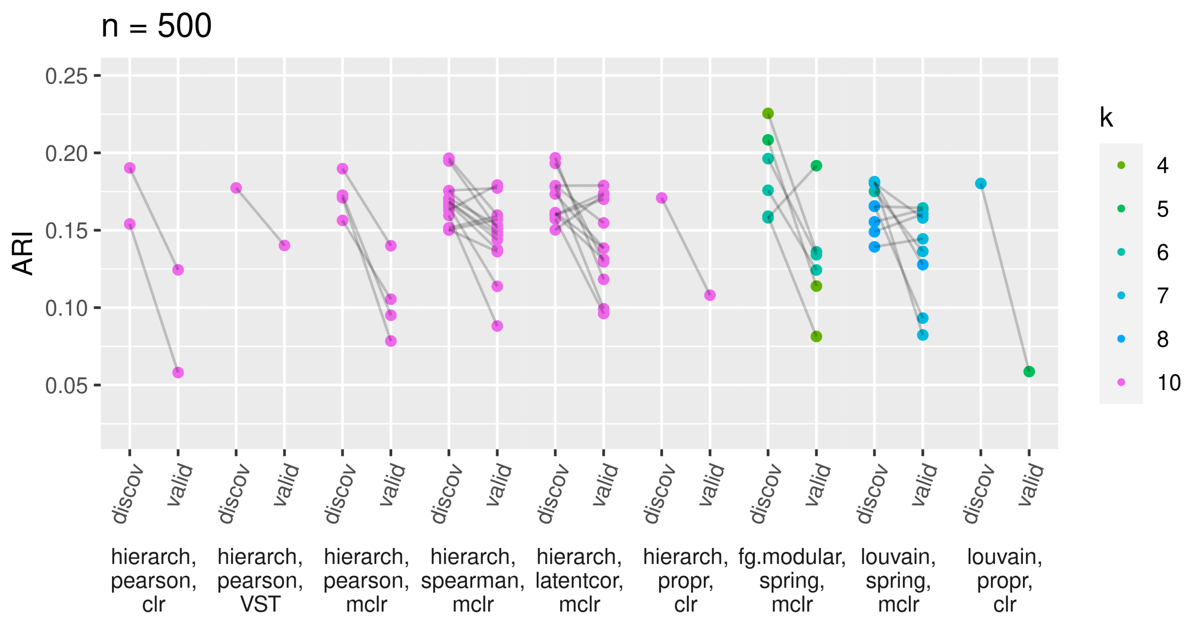


**Fig A.13.** Best ARIs for the clustering of bacterial genera on the discovery data, compared with the results on validation data, $n = 500$

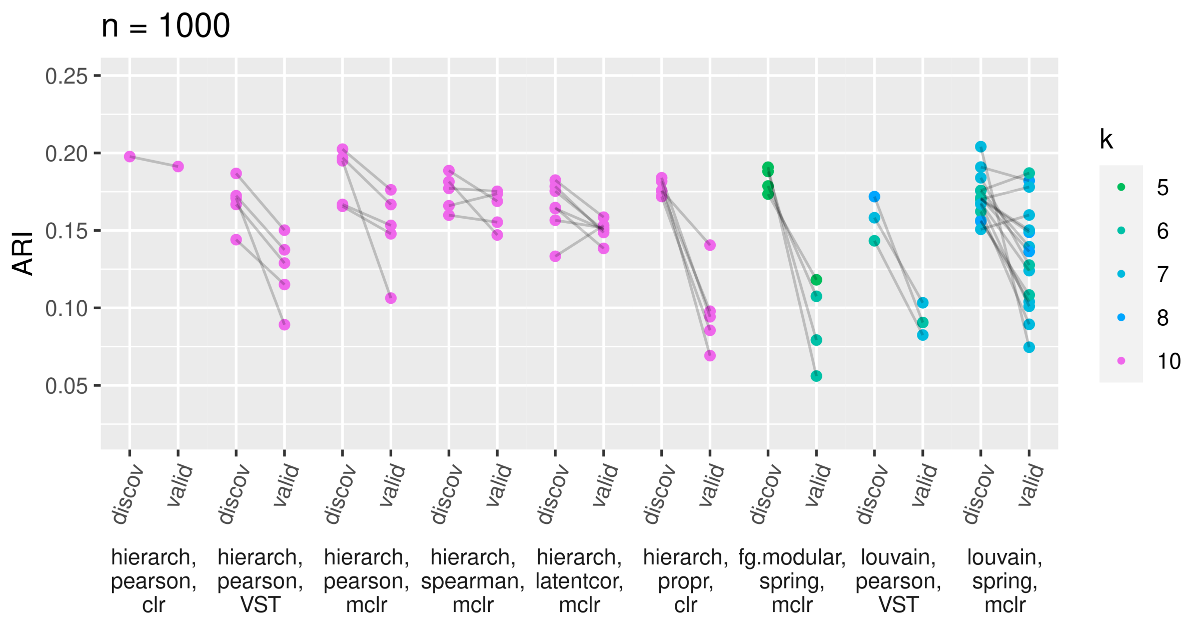


**Fig A.14.** Best ARIs for the clustering of bacterial genera on the discovery data, compared with the results on validation data, $n = 1000$

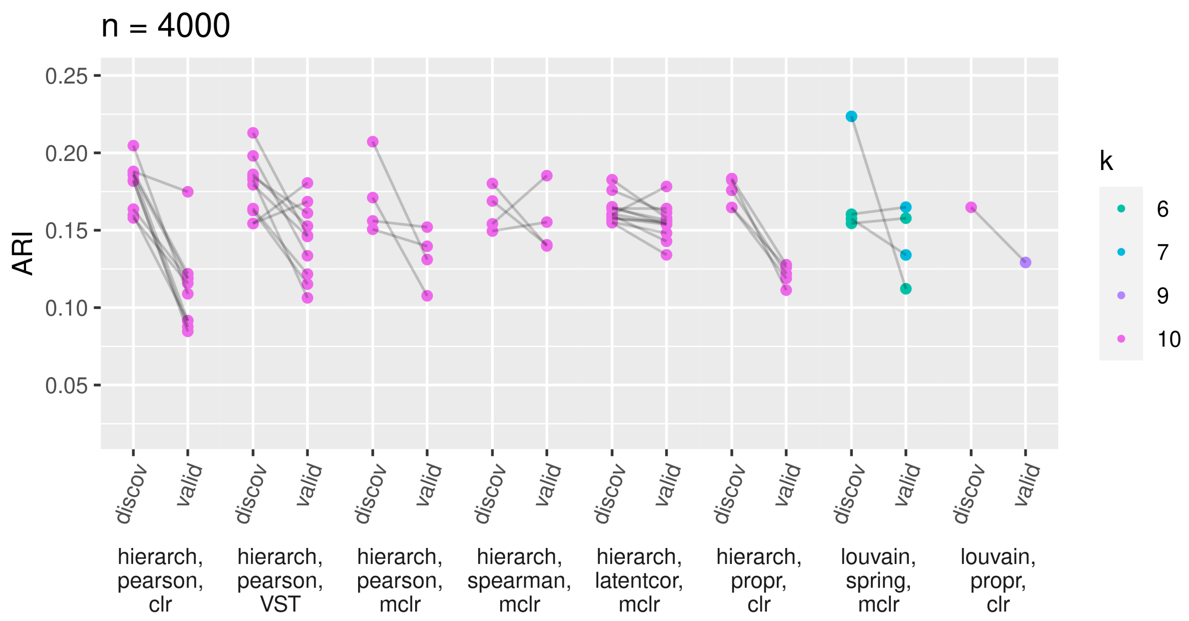


**Fig A.15.** Best ARIs for the clustering of bacterial genera on the discovery data, compared with the results on validation data, $n = 4000$

#### A.2 Research task 2: Hub detection

Fig A.16-A.20 display the results of the hub detection applied to the discovery data. As in the previous section, for each method combination on the $x$-axis, the 50 results obtained from 50 different discovery datasets are summarized as boxplots, indicating the number of detected hubs. Outliers are marked by black crosses. Results that were picked as the "best result" in one of the 50 samplings are marked by red squares.

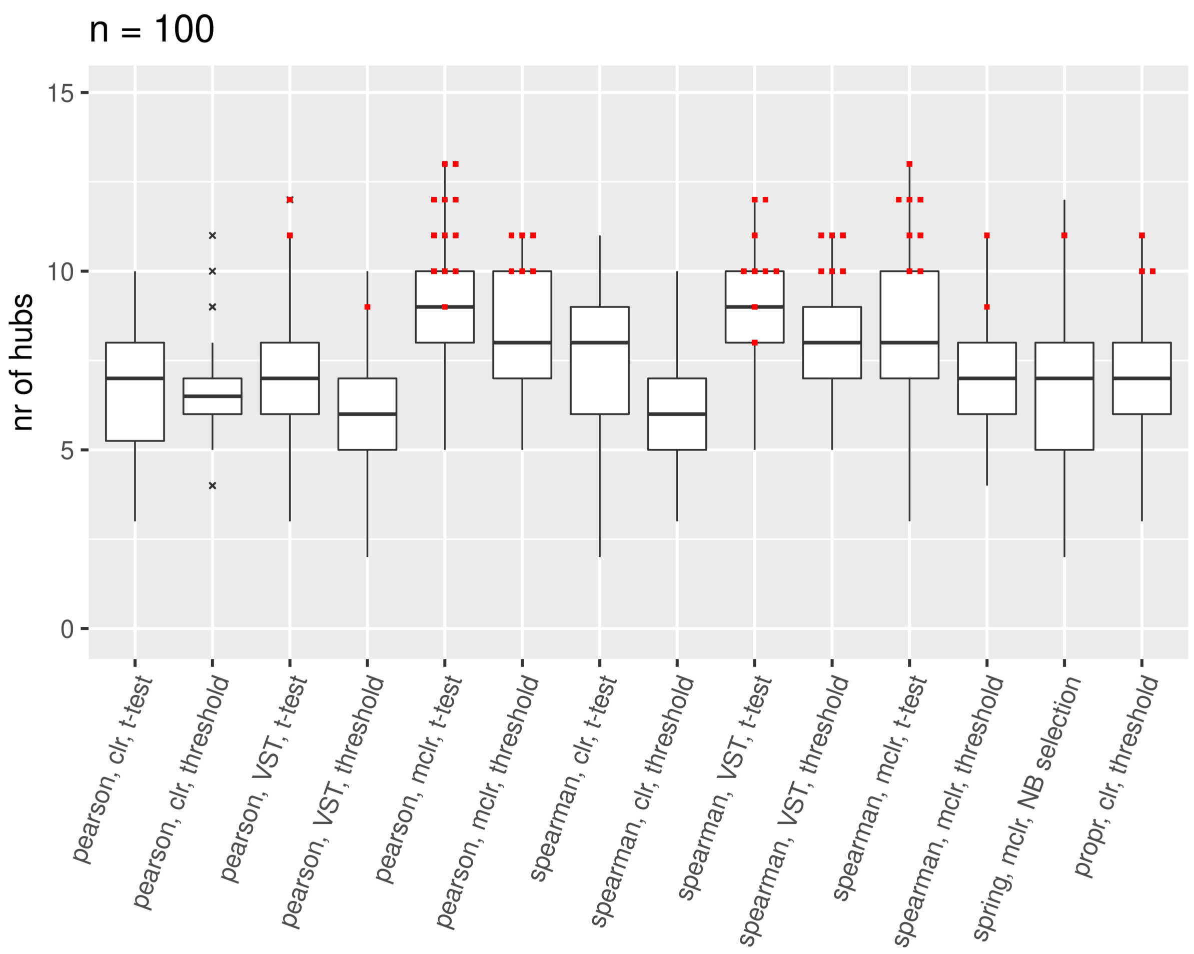


**Fig A.16.** Results for hub detection on the discovery data, $n = 100$

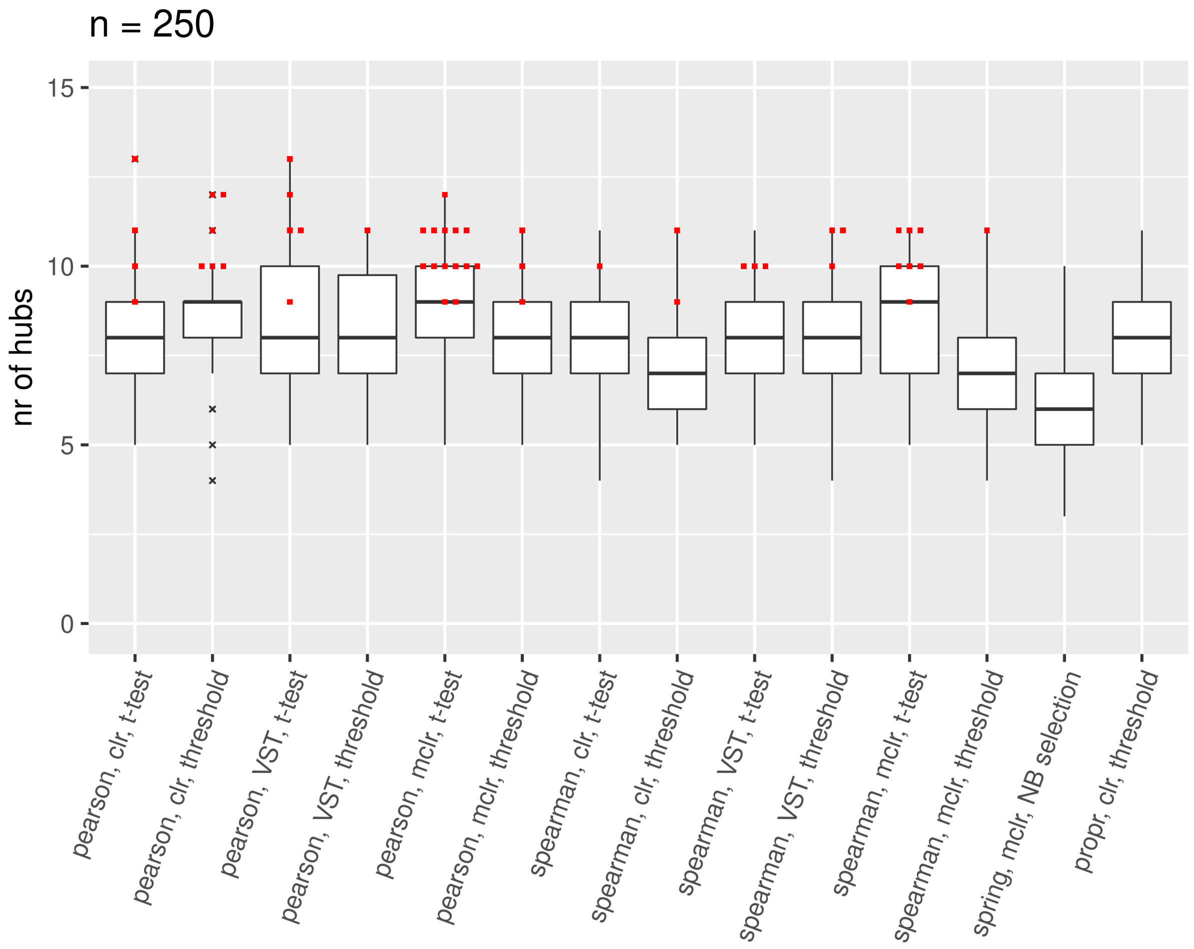


**Fig A.17.** Results for hub detection on the discovery data, $n = 250$

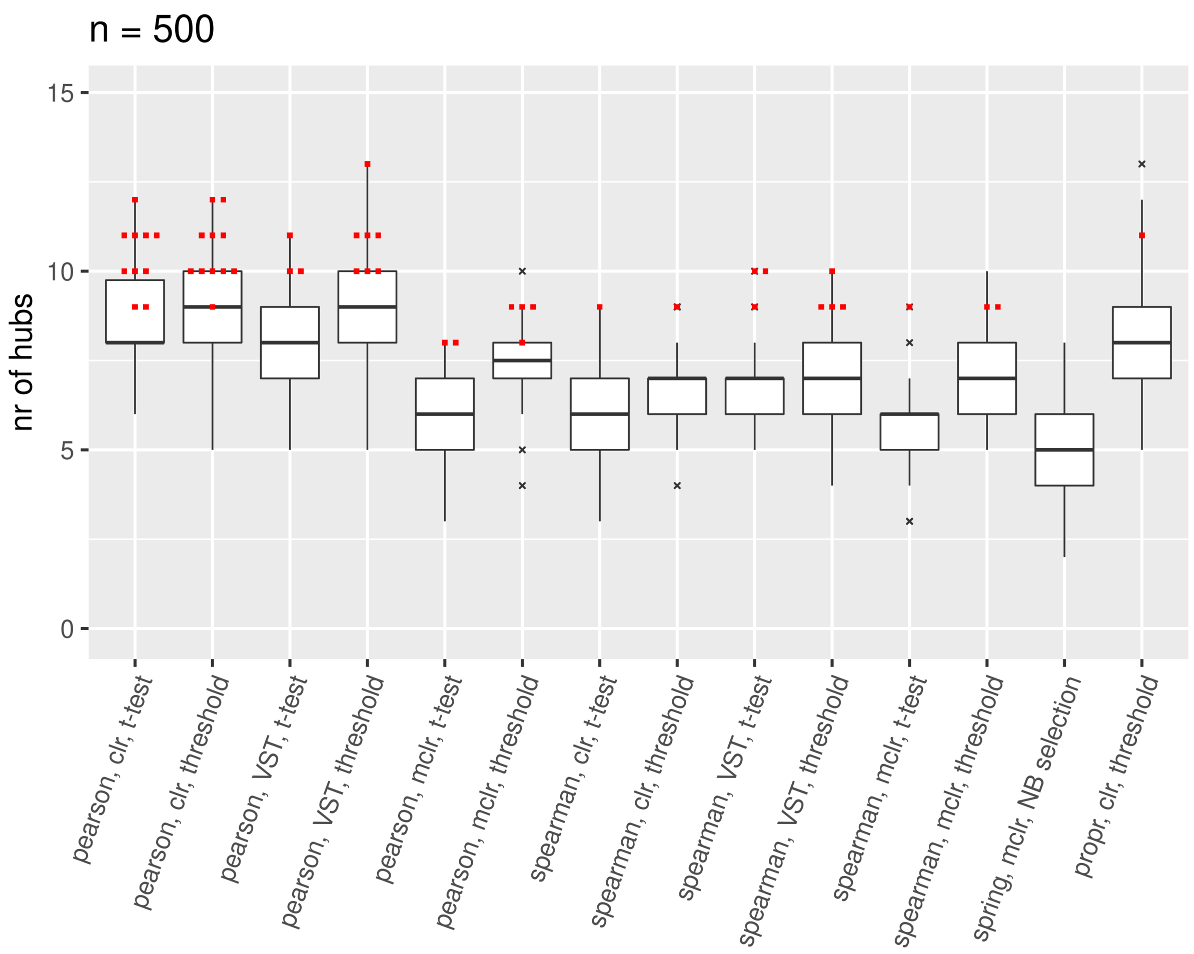


**Fig A.18.** Results for hub detection on the discovery data, $n = 500$

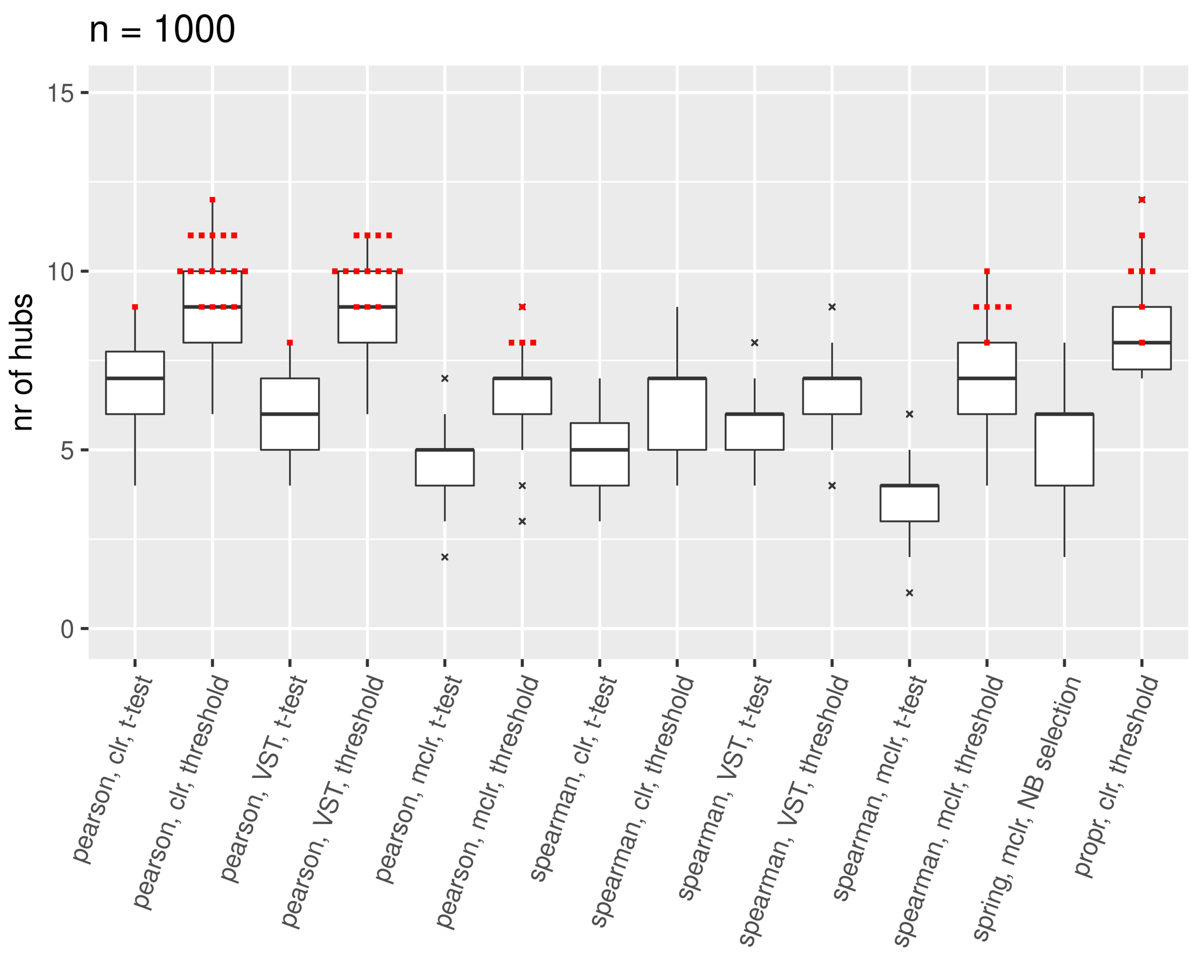


**Fig A.19.** Results for hub detection on the discovery data, $n = 1000$

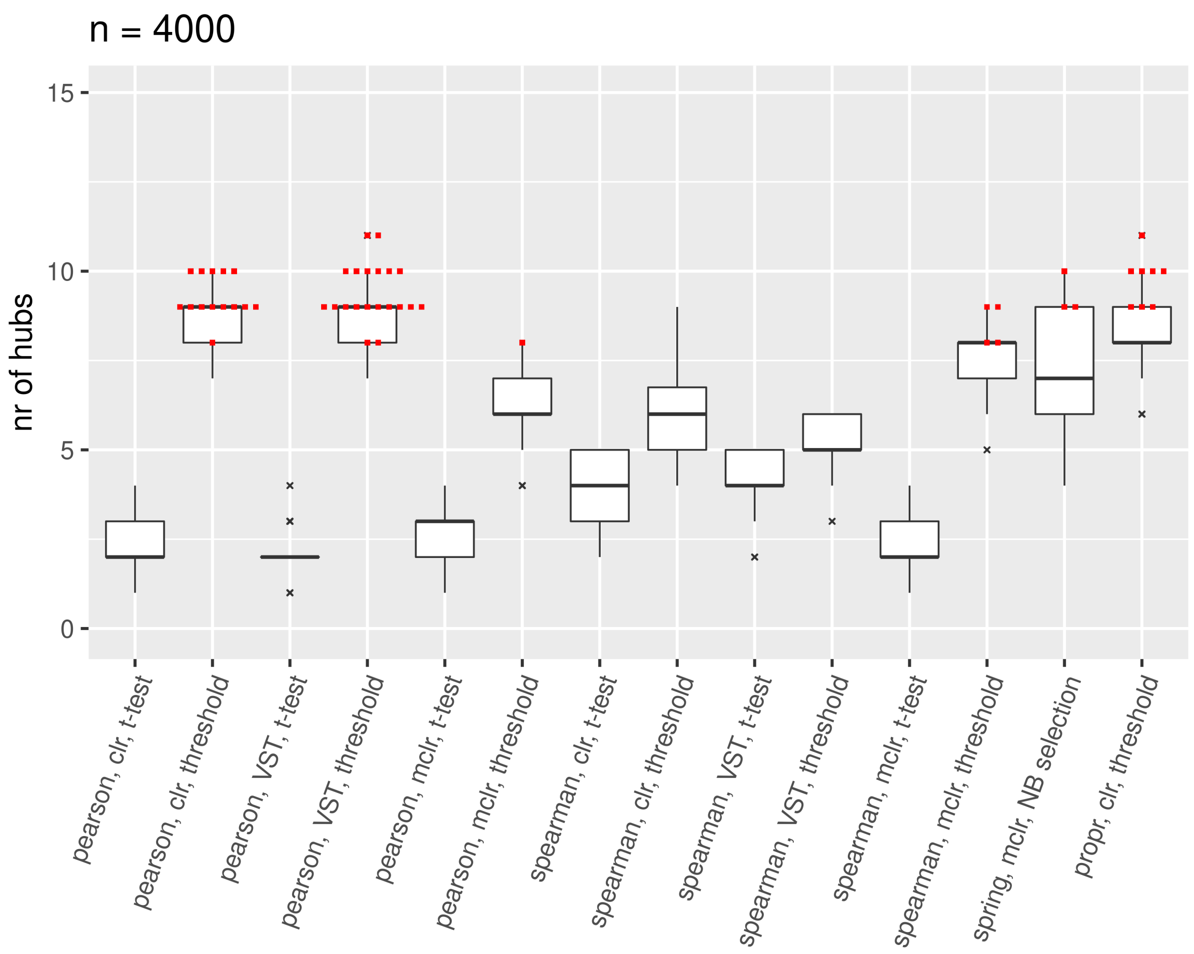


**Fig A.20.** Results for hub detection on the discovery data, $n = 4000$

There is not one single method combination that always yields the highest number of hubs. At $n = 100$, the best results are often found by Pearson correlation with mclr normalization, and Spearman correlation with VST or mclr normalization. With increasing sample size, Pearson correlation with clr or VST normalization frequently yields high number of hubs. As Fig A.19 and Fig A.20 show, for $n = 1000$ and $n = 4000$, sparsification of the network with the $t$-test generally leads to lower number of hubs compared to sparsification with the threshold method. At these sample sizes, the threshold method has a stronger sparsification effect than the $t$-test (given the chosen threshold of 0.15) and sparser networks tend to have more hubs for the chosen hub definition.
We consider the results of applying the chosen method combinations to the validation data. For each method combination that was chosen at least once as the “best” one, Fig A.21-A.24 display the number of hubs obtained by the method on the discovery data vs. the number obtained by the same method on the validation data, where each square-dot combination corresponds to one of the 50 samplings. The results on discovery and validation data are connected by lines.

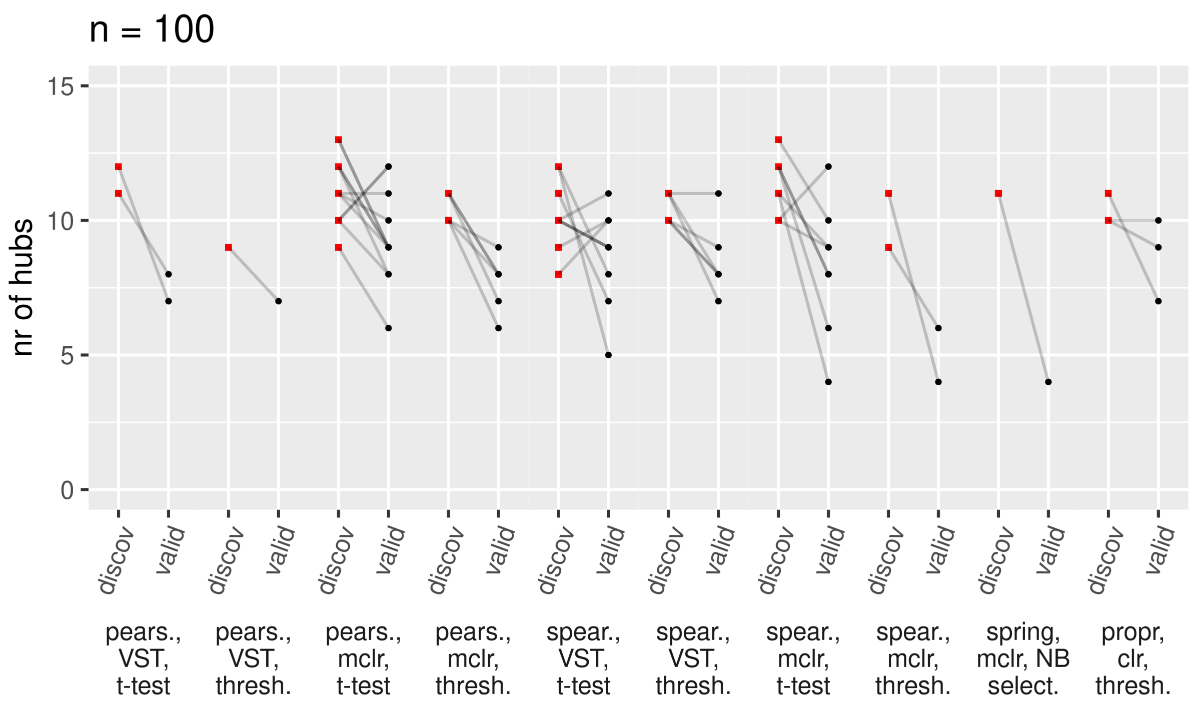


**Fig A.21.** Highest numbers of hubs for the hub detection on the discovery data, compared with the results on validation data, $n = 100$

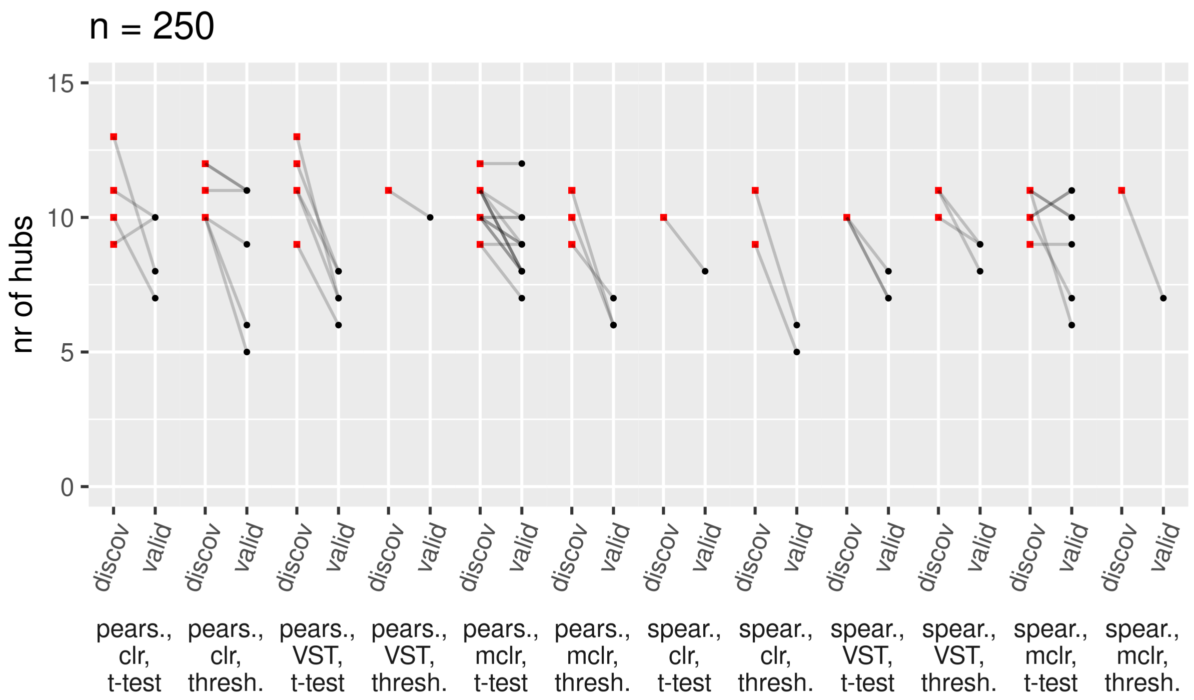


**Fig A.22.** Highest numbers of hubs for the hub detection on the discovery data, compared with the results on validation data, $n = 250$

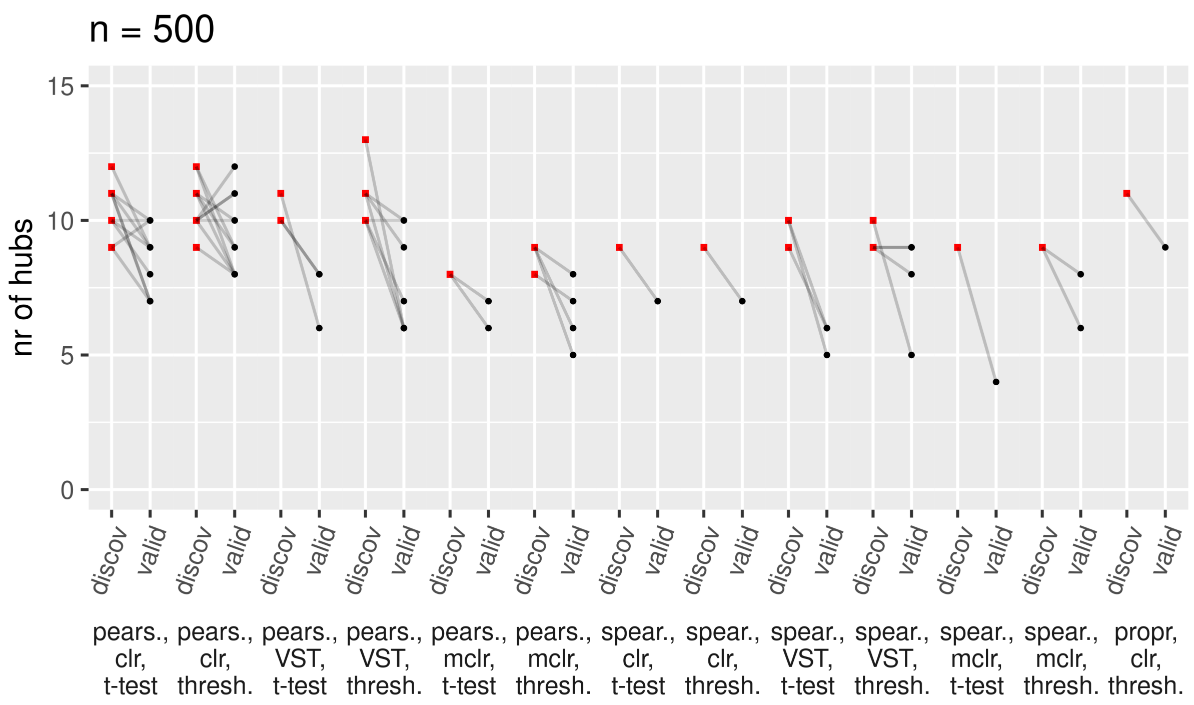


**Fig A.23.** Highest numbers of hubs for the hub detection on the discovery data, compared with the results on validation data, $n = 500$

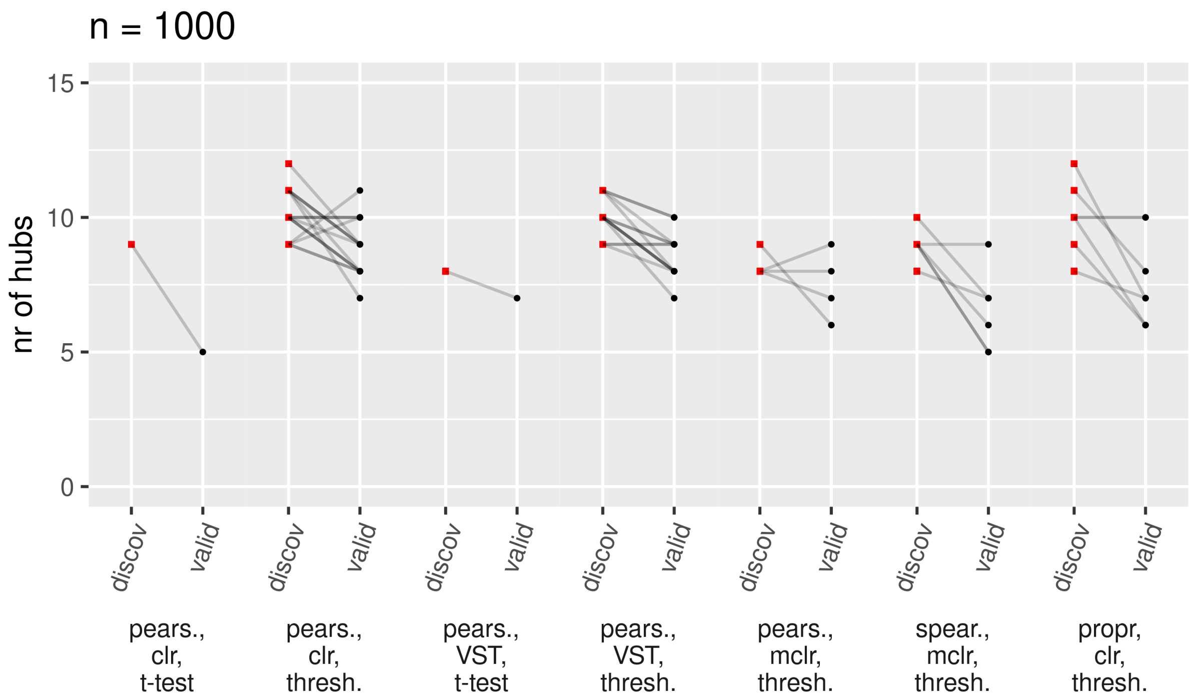


**Fig A.24.** Highest numbers of hubs for the hub detection on the discovery data, compared with the results on validation data, $n = 1000$

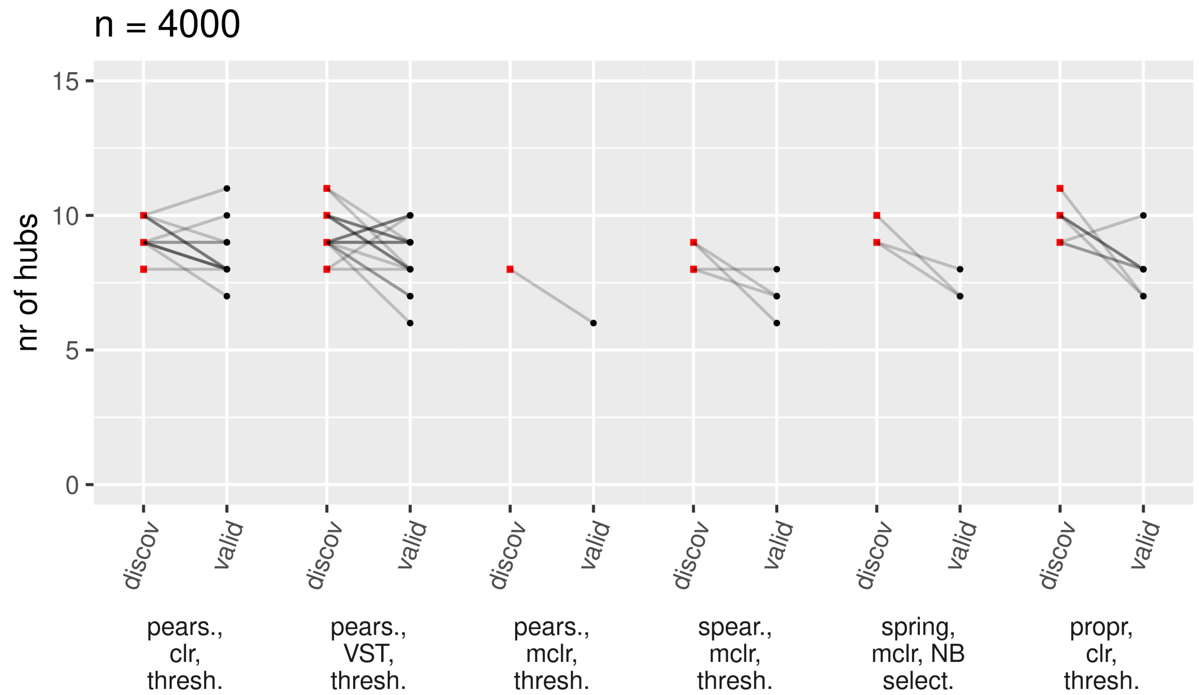


**Fig A.25.** Highest numbers of hubs for the hub detection on the discovery data, compared with the results on validation data, $n = 4000$

The lines point downwards in the majority of the 50 samplings, indicating worse results regarding the network's hubbiness on the validation data.

#### A.3 Research task 3: Differential network analysis

In the same format as in the previous section, Fig A.26-A.28 show the results of the differential network analysis on the discovery data over 50 samplings. Boxplots summarize the GCDs between the microbial network based on the non-antibiotics samples vs. the network based on the antibiotics samples. GCDs that were picked as the "best" results are marked by red squares.

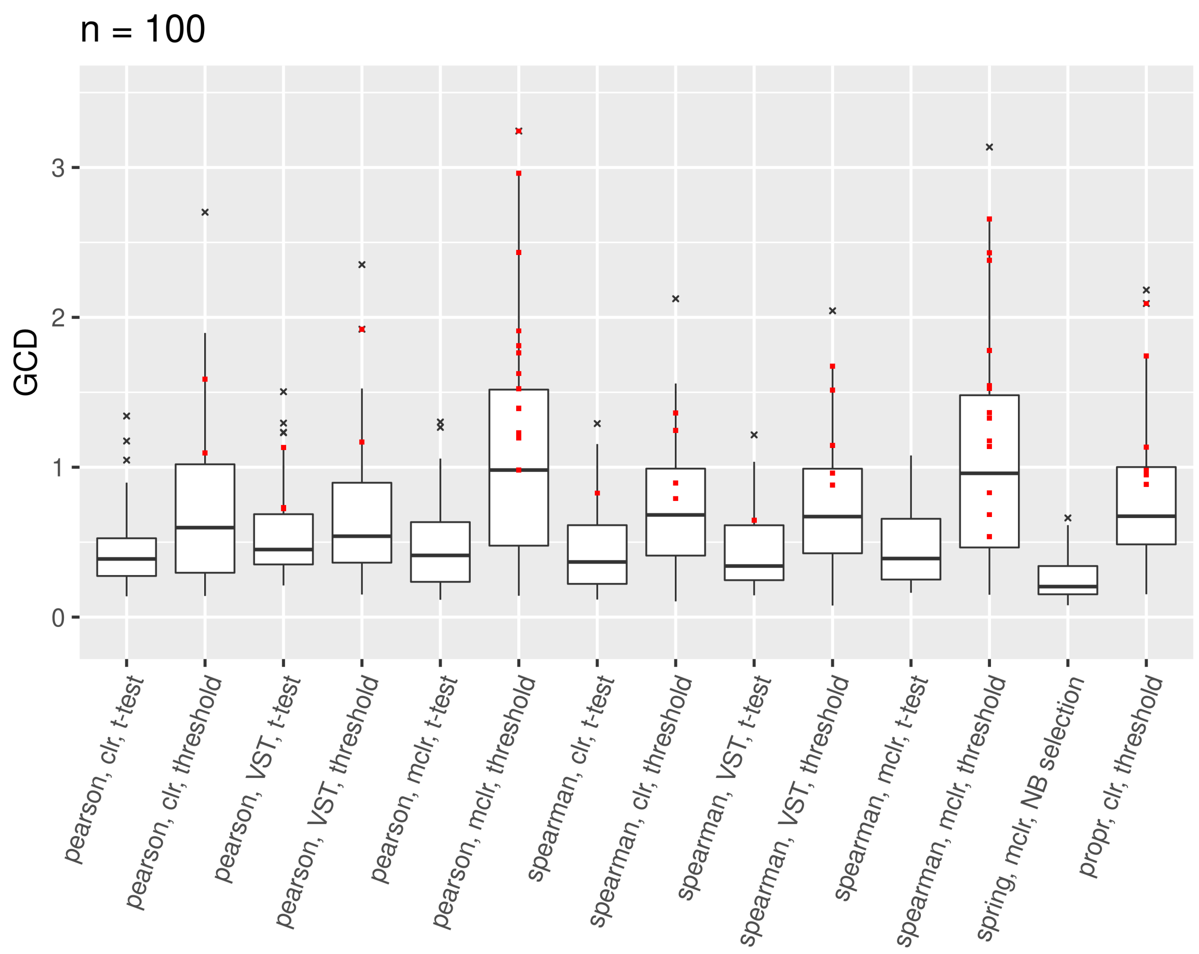


**Fig A.26.** Results for differential network analysis on the discovery data, $n = 100$

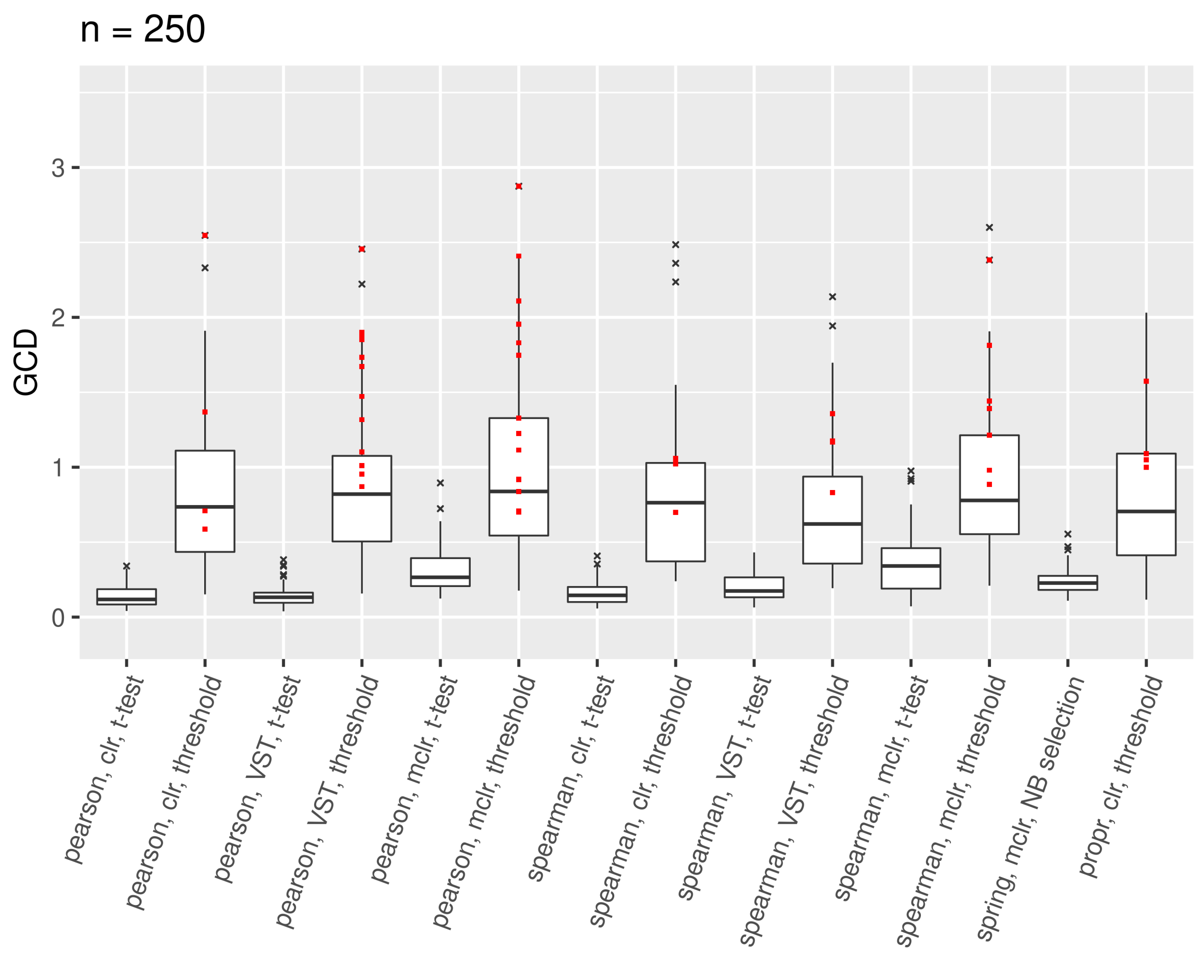


**Fig A.27.** Results for differential network analysis on the discovery data, $n = 250$

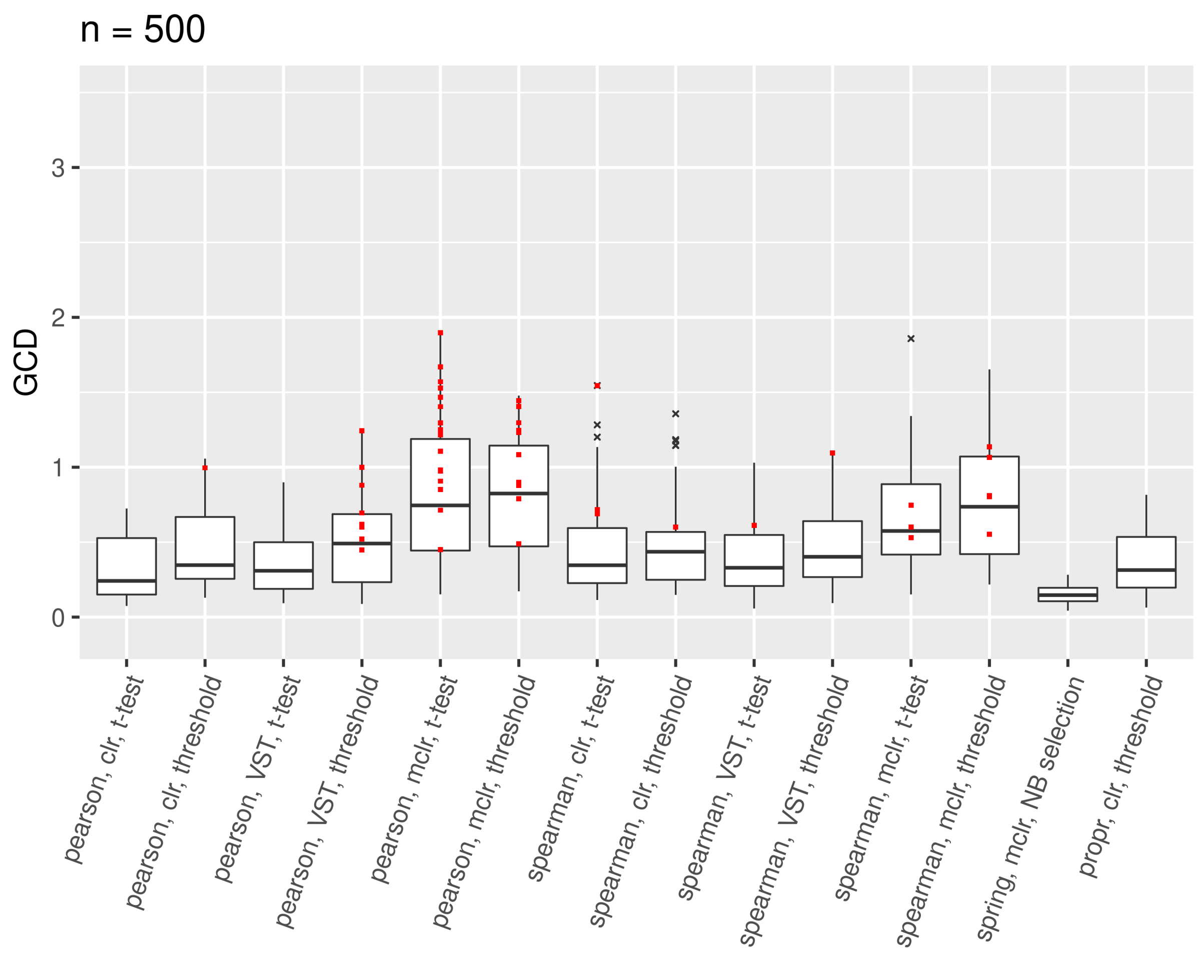


**Fig A.28.** Results for differential network analysis on the discovery data, $n = 500$

Similar to hub detection, there is no superior method combination that always leads to best results, i.e., highest GCD values. Notably, sparsification via $t$-test never leads to best results for $n = 100$ and $n = 250$, but only for $n = 500$. However, a general trend cannot be confirmed due to the limited sample size in this research task.
Fig A.29-A.31 show the results of applying the best method combinations to the validation data and compare these to the results on the discovery data. Over-optimism is indicated by downward lines, which is the case in about 75% of the 50 samplings.

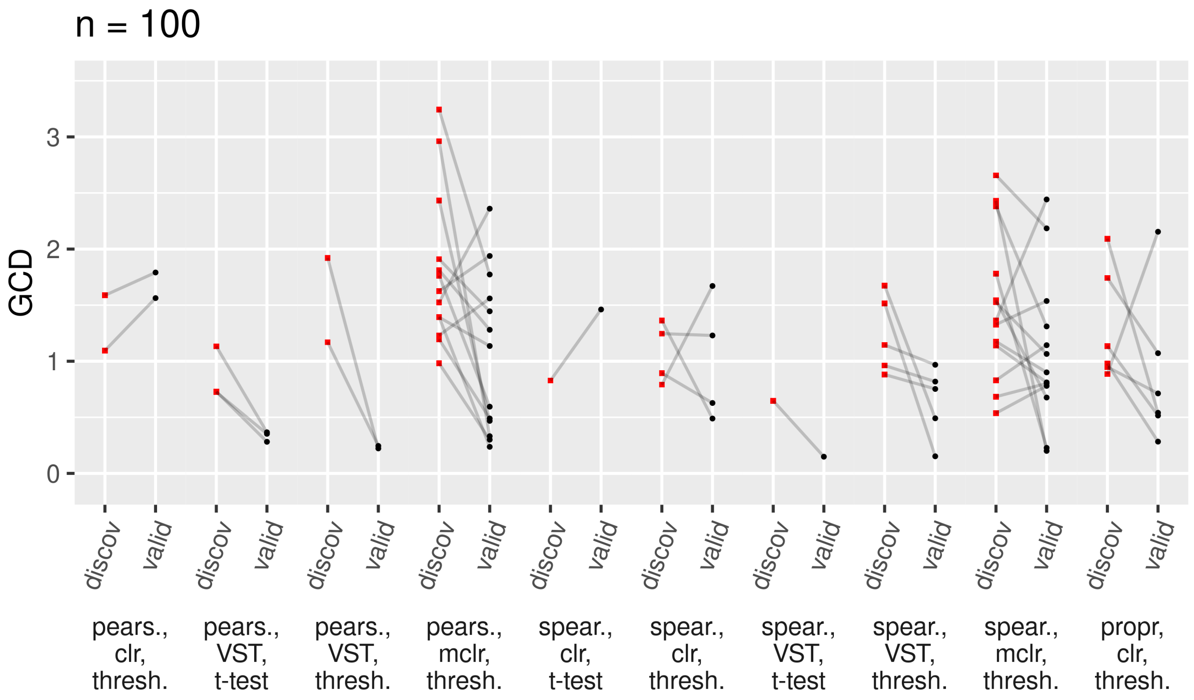


**Fig A.29.** Largest GCDs for the differential network analysis on the discovery data, compared with the results on validation data, $n = 100$

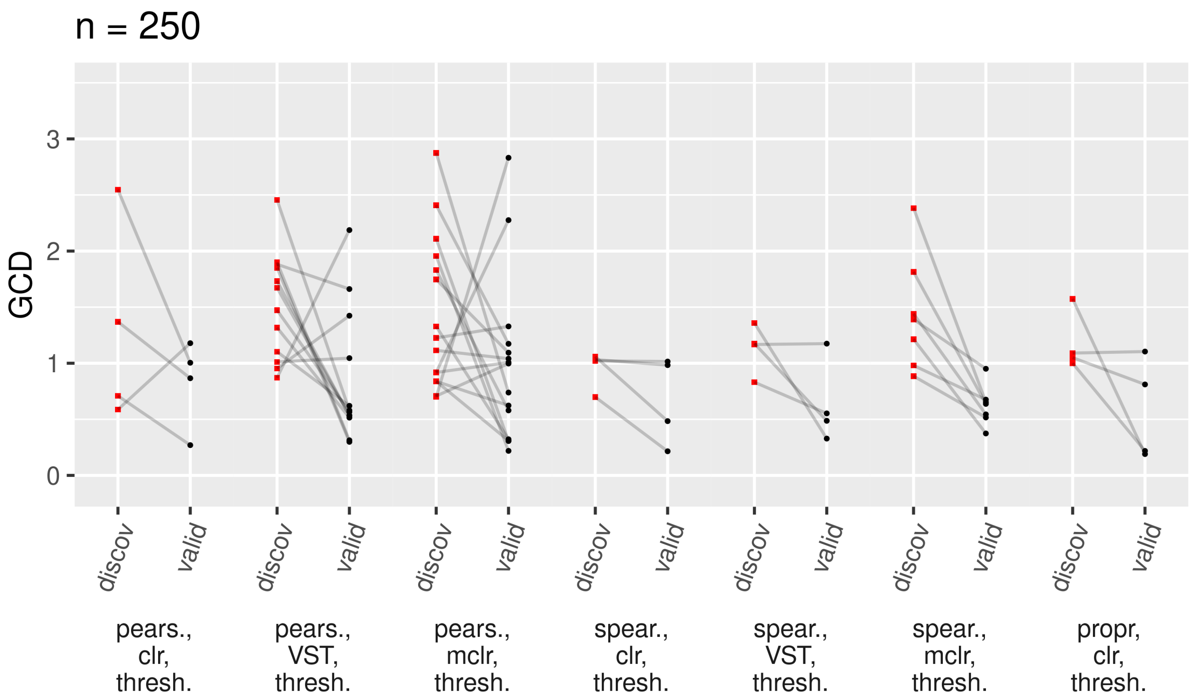


**Fig A.30.** Largest GCDs for the differential network analysis on the discovery data, compared with the results on validation data, $n = 250$

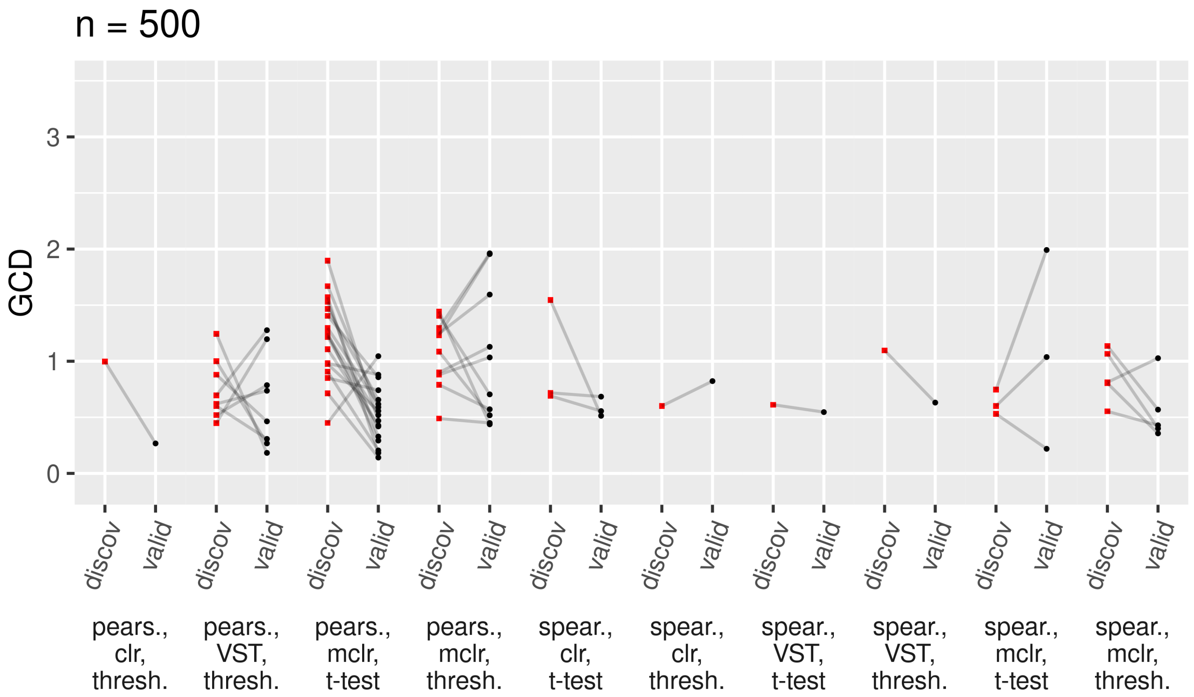


**Fig A.31.** Largest GCDs for the differential network analysis on the discovery data, compared with the results on validation data, $n = 500$
